# Hydrogen-based metabolism – an ancestral trait in lineages sibling to the Cyanobacteria

**DOI:** 10.1101/328856

**Authors:** Paula B. Matheus Carnevali, Frederik Schulz, Cindy J. Castelle, Rose Kantor, Patrick Shih, Itai Sharon, Joanne M. Santini, Matthew Olm, Yuki Amano, Brian C. Thomas, Karthik Anantharaman, David Burstein, Eric D. Becraft, Ramunas Stepanauskas, Tanja Woyke, Jillian F. Banfield

**Affiliations:** Department of Earth and Planetary Science, University of California, Berkeley, CA, USA.; DOE Joint Genome Institute, Walnut Creek, CA, USA.; Feedstocks Division, Joint BioEnergy Institute, Emeryville, CA, USA.; Environmental Genomics and Systems Biology Division, Lawrence Berkeley National Laboratory, Berkeley, CA, USA.; Institute of Structural & Molecular Biology, Division of Biosciences, University College London, London, UK.; Department of Plant and Microbial Biology, University of California, Berkeley, CA, USA.; Nuclear Fuel Cycle Engineering Laboratories, Japan Atomic Energy Agency, Tokai, Ibaraki, Japan.; Horonobe Underground Research Center, Japan Atomic Energy Agency, Horonobe, Hokkaido, Japan.; Bigelow Laboratory for Ocean Sciences, East Boothbay, ME, USA.; Earth Sciences Division, Lawrence Berkeley National Laboratory, Berkeley, California, USA.; Chan Zuckerberg Biohub, San Francisco, CA, USA.; Innovative Genomics Institute, Berkley, CA, USA.; Department of Civil and Environmental Engineering, University of California, Berkeley, CA, USA.; Department of Plant Biology, University of California, Davis, Davis, CA, USA.; Migal Galilee Research Institute, Kiryat Shmona, 11016, Israel.; Tel Hai College, Upper Galilee 12210, Israel.; Department of Bacteriology, University of Wisconsin-Madison, Madison, WI, USA.; School of Molecular and Cell Biology and Biotechnology, George S. Wise Faculty of Life Sciences, Tel Aviv University, Tel Aviv 69978, Israel.

## Abstract

The metabolic machinery from which microbial aerobic respiration evolved is tightly linked to the origins of oxygenic Cyanobacteria (Oxyphotobacteria). Even though the majority of Oxyphotobacteria are photoautotrophs and can use carbohydrates with oxygen (O_2_) as the electron acceptor, all are fermenters under dark anoxic conditions. Studies suggest that the ancestor of Oxyphotobacteria may have used hydrogen (H_2_) as an electron donor and that two types of NiFe hydrogenases are essential for its oxidation. Melainabacteria and Sericytochromatia, close phylogenetic neighbors to Oxyphotobacteria comprise fermentative and aerobic representatives, or organisms capable of both. Margulisbacteria (candidate divisions RBX-1 and ZB3) and Saganbacteria (candidate division WOR-1), a novel cluster of bacteria phylogenetically related to Melainabacteria, Sericytochromatia and Oxyphotobacteria may further constrain the metabolic platform in which oxygenic photosynthesis and aerobic respiration arose. Here, we predict the metabolisms of Margulisbacteria and Saganbacteria from new and published metagenome-assembled genomes (MAGs) and single amplified genomes (SAGs), and compare them to their phylogenetic neighbors. Sediment-associated Margulisbacteria are predicted to have a fermentation-based metabolism featuring a variety of hydrogenases, a nitrogenase for nitrogen (N_2_) fixation, and electron bifurcating complexes involved in cycling of ferredoxin and NAD(P)H. Overall, the genomic features suggest the capacity for metabolic fine-tuning under strictly anoxic conditions. In contrast, the genomes of Margulisbacteria from the ocean ecosystem encode an electron transport chain that supports aerobic growth. Similarly, some Saganbacteria genomes encode various hydrogenases, and others may have the ability to use O2 under certain conditions *via* a putative novel type of heme copper O2 reductase. Like Melainabacteria and Sericytochromatia, Margulisbacteria and Saganbacteria have diverse energy metabolisms capable of fermentation, and aerobic or anaerobic respiration. In summary, our findings support the hypothesis that the ancestor of these groups was an anaerobe in which fermentation and H_2_ metabolism were central metabolic features. Our genomic data also suggests that contemporary lineages sibling to the Oxyphotobacteria may have acquired the ability to use O_2_ as a terminal electron acceptor under certain environmental conditions.

## Introduction

Oxygenic photosynthesis evolved in Cyanobacteria, now referred to as Oxyphotobacteria (Soo et al., 2014; Shih et al., 2017; Soo et al., 2017), and ultimately led to the Great Oxidation Event (GOE) ∼ 2.3 billion years ago (Luo et al., 2016). Microorganisms had access to oxygen (O_2_) as an electron acceptor following the GOE, and thus higher energetic yield from respiratory processes. However, use of O_2_ required the evolution of redox complexes in the electron transport chain (ETC). These innovations may have occurred in ancestral Oxyphotobacteria that immediately benefited from the opportunities presented by O_2_ availability.

The metabolic potential of lineages phylogenetically related to Oxyphotobacteria is of great interest from the perspective of evolution and co-evolution of life on this planet (Soo et al., 2014; Shih et al., 2017; Soo et al., 2017). Recently, several publications have investigated the biology of Melainabacteria and Sericytochromatia, groups that branch adjacent to Oxyphotobacteria and are represented by non-photosynthetic organisms (Di Rienzi et al., 2013; Soo et al., 2014; Soo et al., 2017). It has been suggested that the machinery for aerobic respiration was acquired independently by Sericytochromatia (Soo et al., 2017), and like Oxyphotobacteria, some members of the Sericytochromatia and Melainabacteria have ETCs. The most simple ETC is composed of (i) an electron entry point, usually a dehydrogenase that oxidizes reduced electron carriers (*e.g.*, nicotinamide adenine nucleotide – NADH and succinate), (ii) membrane electron carriers (*e.g.,* quinones), and (iii) an electron exit point such a heme-copper oxygen reductase or a cytochrome *bd* oxidase. However, if there are periplasmic electron donors, an intermediary quinol:electron acceptor oxidoreductase complex may also be involved (Refojo et al., 2012). Both the electron entry and exit protein complexes often act as H^+^ pumps, contributing to the creation of a proton motive force (PMF; or electrochemical potential) that can be used for energy generation via the ATP synthase.

Oxyphotobacteria use light as the source of energy to establish an electrochemical H^+^ potential during photosynthetic electron transport and also conserve energy from reducing a high potential terminal electron acceptor (O_2_) during aerobic respiration. Aerobic Sericytochromatia may use chemical energy rather than light to generate an electrochemical potential gradient (Soo et al., 2017). During evolution, these high electrode potential respiratory chains may have replaced fermentation-linked complexes responsible for ion translocation across membranes in more ancient organisms. Sequential translocation of ions across membranes is required in fermentation-based metabolisms as the energy yield per translocation is low (Boyd et al., 2014). Analysis of new bacterial genomes from sibling groups to the Oxyphotobacteria may provide information about their ancestral energy metabolism and thus constrain the metabolic platform into which complex ETCs evolved.

Here we predict the metabolic capacities of bacteria from lineages that are sibling to Oxyphotobacteria, Melainabacteria, and Sericytochromatia. Genomes for members of the Margulisbacteria (RBX-1) and Saganbacteria (WOR-1) were previously reconstructed from estuarine sediments and groundwater (Baker et al., 2015; Anantharaman et al., 2016; Probst et al., 2018). For Margulisbacteria, here we describe two groups that we refer to as Riflemargulisbacteria (given the derivation of draft genome sequences from the Rifle research site) (Anantharaman et al., 2016), and four groups we refer to as Marinamargulisbacteria (given the derivation of the single-cell draft genomes from ocean sites). Marinamargulisbacteria was first observed in a clone library of SSU rRNA genes from Zodletone spring (Oklahoma) and identified as candidate division ZB3 (Elshahed et al., 2003). In this study, we predict how organisms from these lineages conserve energy, identify their unique strategies for reoxidation of reducing equivalents, and propose the interconnectedness between H_2_ and O_2_ metabolism. We found that H2 metabolism is a common feature of the modern groups that cannot aerobically respire aerobically, and suggest that hydrogenases may have been central to the lifestyles of the ancestors of Oxyphotobacteria, Melainabacteria, Sericytochromatia, Margulisbacteria and Saganbacteria.

## Results

### Identification of representative genomes from each lineage

In this study, we analyzed four publicly available and eight newly reconstructed genomes of Margulisbacteria and propose two distinct clades within this lineage: Riflemargulisbacteria (two groups) and Marinamargulisbacteria (four groups). These clades were defined using metagenome-assembled genomes (MAGs) of bacteria from the sediments of an aquifer adjacent to the Colorado River, Rifle, USA and single-cell amplified genomes (SAGs) from four ocean environments (Supplementary Table 1). We also analyzed 26 publicly available MAGs of Saganbacteria from groundwater with variable O_2_ concentrations (Anantharaman et al., 2016), of which four genomes are circularized. In addition we analyzed eight publicly available MAGs of Melainabacteria (Anantharaman et al., 2016), and five new Melainabacteria genomes that were reconstructed from metagenomic datasets for microbial communities sampled from human gut and groundwater (Supplementary Table 1). We identified representative genomes for groups of genomes that share 95.0 − 99.0% average nucleotide identity (ANI) (Supplementary Table 1) and focus our discussion on these genomes. Metabolic predictions for genomes that were estimated to be medium or high quality (Bowers et al., 2017) (Supplementary Table 1) were established using gene annotations and confirmed using Hidden Markov Models built from the KEGG database (Supplementary Table 2). Key aspects of energy metabolism predicted for Riflemargulisbacteria, Marinamargulisbacteria, Saganbacteria and Melainabacteria were compared. Phylogenetic analyses of genes encoding key metabolic functions also included publicly available genomes for Sericytochromatia (Soo et al., 2017), newly reported genomes for relatives of Marinamargulisbacteria (Parks et al., 2017), and other reference genomes.

### Margulisbacteria and Saganbacteria are the closest relatives of Oxyphotobacteria, Melainabacteria and Sericytochromatia

In our phylogenetic analysis, Margulisbacteria and Saganbacteria represent two major lineages that are sibling to Sericytochromatia, Melainabacteria and Oxyphotobacteria (Fig. 1a, 1b). Together, Margulisbacteria and Saganbacteria are positioned basal to the Oxyphotobacteria. Importantly, the affiliation of these groups with the Oxyphotobacteria is consistent for phylogenetic trees based on 56 universal single copy proteins (Yu et al., 2017) and 16 ribosomal proteins (Hug et al., 2016) (Fig. 1a, Supplementary Fig. 1, 2). However, the precise position of these three lineages differs depending on the set of markers and phylogenetic models used for tree construction (Fig. 1a, Supplementary Fig. 2) and the deep branching order often cannot be well resolved (Hug et al., 2016; Parks et al., 2017). In addition, the position of Margulisbacteria, Saganbacteria, and Oxyphotobacteria relative to the Thermoanaerobacterales (Fig. 1a) will require further investigation once more genomes for these groups become available. Similar to Sericytochromatia, Oxyphotobacteria and Melainabacteria, the clustering of Saganbacteria, Riflemargulisbacteria and Marinamargulisbacteria is well supported (posterior probability > 0.97). However, the analysis of shared protein families among lineages, suggested that Riflemargulisbacteria, Marinamargulisbacteria, and Saganbacteria share less protein families than Oxyphotobacteria, Melainabacteria and Sericytochromatia do (Fig. 1c). Despite the closer phylogenetic relationship between Riflemargulisbacteria and Marinamargulisbacteria, Riflemargulisbacteria share a greater proportion of protein families with Saganbacteria than with Marinamargulisbacteria, whereas Marinamargulisbacteria share most protein families with the more distantly related Oxyphotobacteria. Taken together, these findings may indicate niche adaptation and a divergent lifestyle of Riflemargulisbacteria and Marinamargulisbacteria.

**Figure 1:**
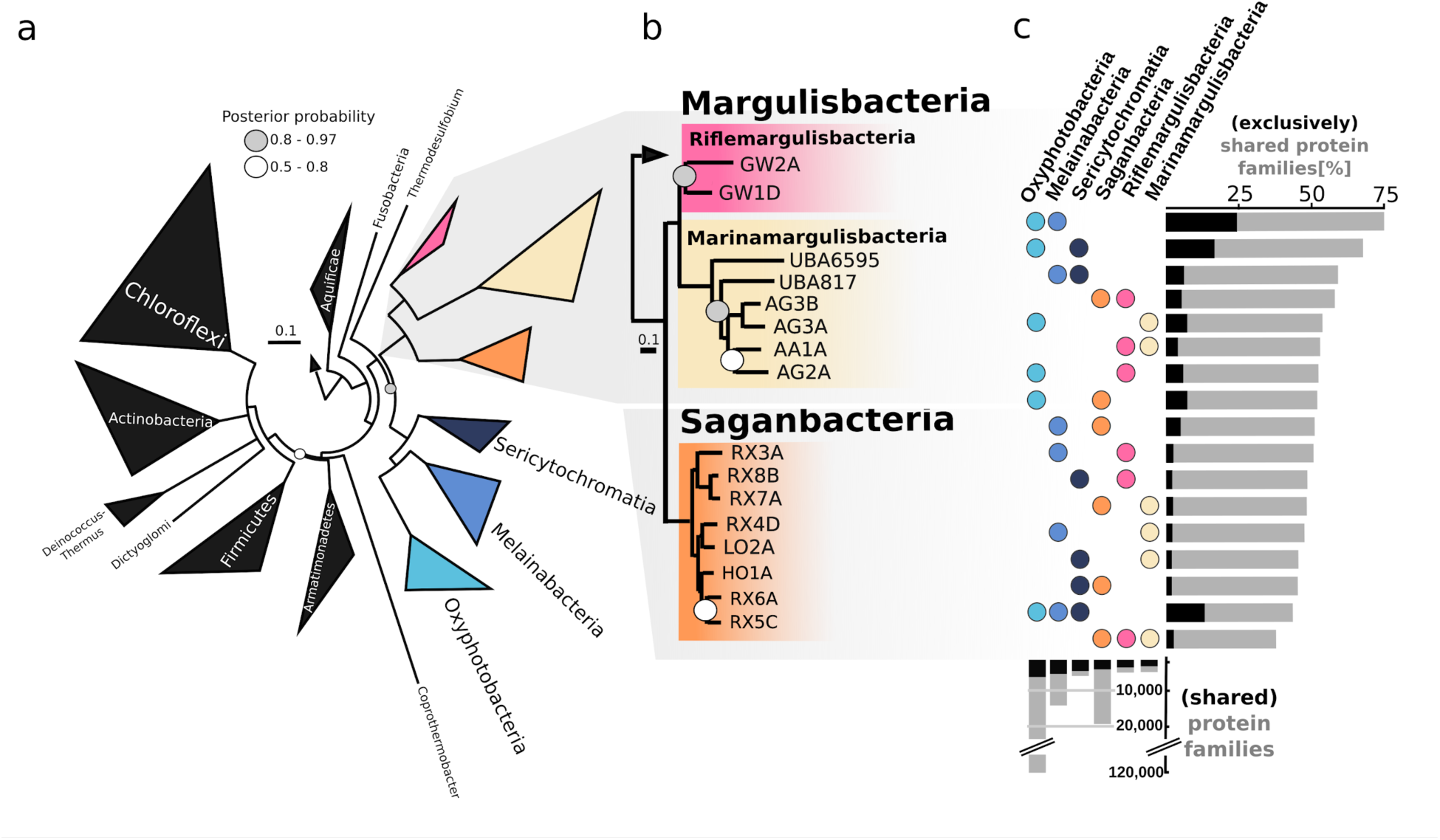
Phylogenetic tree based on fifty-six concatenated proteins and shared protein families in Cyanobacteria (Oxyphotobacteria), Sericytochromatia, Margulisbacteria, Saganbacteria, and Melainabacteria. **a.** Location of Margulisbacteria and Saganbacteria in a de-replicated species tree of the Terrabacteria (detailed tree shown as Supplementary Fig. 1). **b.** Phylogenetic relationship of members of the Margulisbacteria and Saganbacteria. See Supplementary Table 1 for genome identification details **c.** Shared protein families among lineages: rows cover the total number of protein families shared among lineages and dots indicate which lineages are being compared. For example, the top bar indicates that in average 75% of protein families are shared between Oxyphotobacteria and Melainabacteria, out of which 25% are exclusively shared between them. The other 50% includes proteins that are shared with at least one other lineage. At the bottom, bars show the total number of protein families found in a given lineage in gray, and the proportion that is shared with one or more other lineages in black. For details on taxon sampling, tree inference and shared protein families, see Materials and methods.

### Sediment-associated Riflemargulisbacteria: free-living, hydrogen-driven anaerobic heterotrophs

Metabolic analyses based on a representative genome of the sediment-associated Riflemargulisbacteria, Margulisbacteria RA1A (96% completeness, Supplementary Table 1), suggest that members of Riflemargulisbacteria are heterotrophic organisms with a fermentation-based metabolism (Fig. 2). They use hydrogenases and other electron bifurcating complexes to balance their reducing equivalents (NADH and ferredoxin), and rely on membrane-bound protein complexes to generate a proton/sodim (H^+^/Na^+^) potential and to make ATP (Fig. 3a).

**Figure 2:**
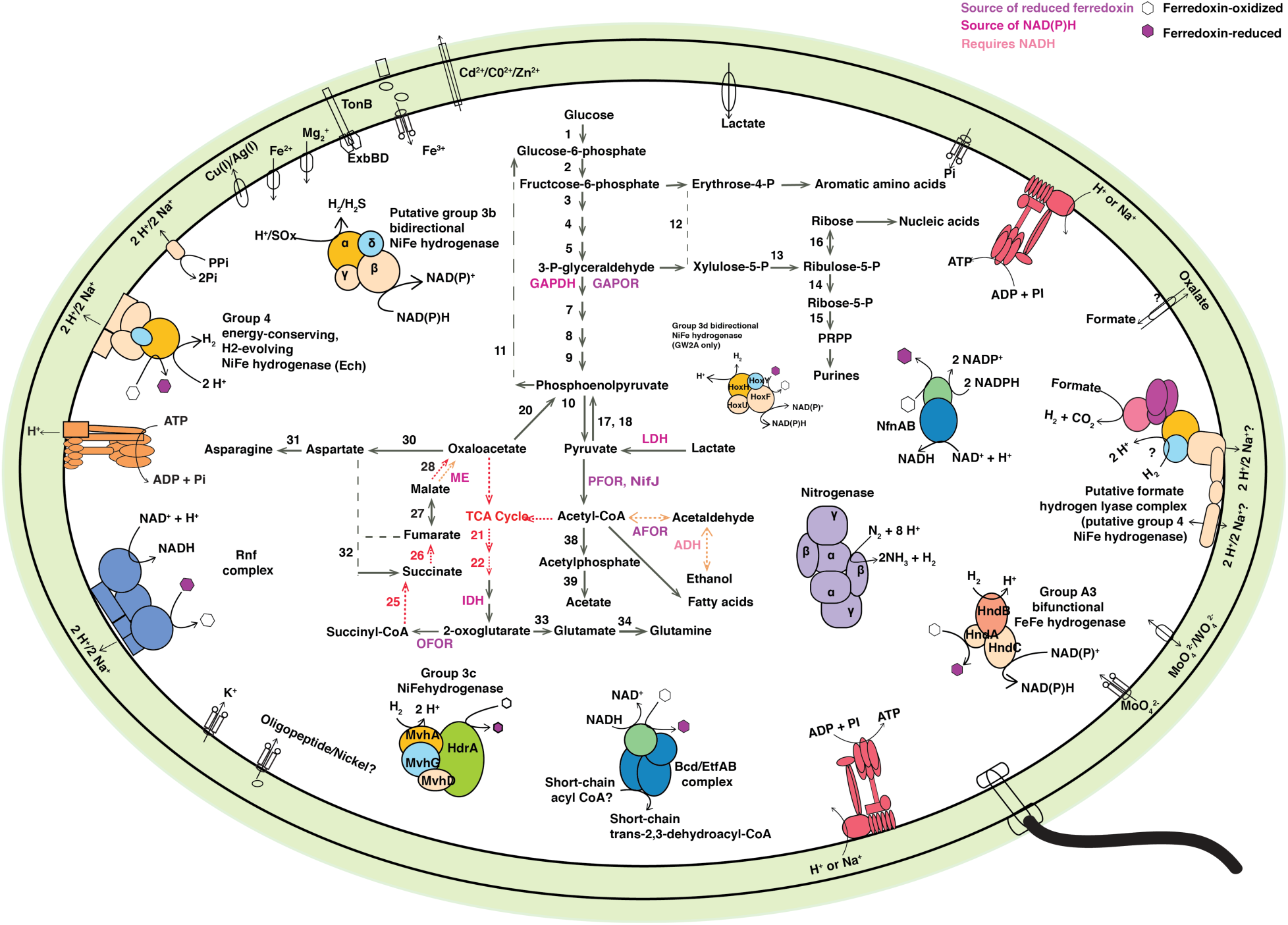
Margulisbacteria RA1A cell cartoon. Key enzymes predicted to be involved in core metabolic pathways include: (1) glucokinase, (2) glucose-6-phosphate isomerase, (3) 6-phosphofructokinase 1, (4) fructose-bisphosphate aldolase class I and II, (5) triosephosphate isomerase, GAPDH: glyceraldehyde 3-phosphate dehydrogenase, GAPOR: glyceraldehyde-3-phosphate dehydrogenase (ferredoxin), (7) phosphoglycerate kinase, (8) phosphoglycerate mutase 2,3-bisphosphoglycerate-dependent and 2,3-bisphosphoglycerate-independent, (9) enolase, (10) pyruvate kinase, (11) fructose-1,6-bisphosphatase III, (12) transketolase, (13) ribulose-phosphate 3-epimerase, (14) ribose 5-phosphate isomerase B, (15) ribose-phosphate pyrophosphokinase, (16) ribokinase, (17) pyruvate, orthophosphate dikinase, (18) pyruvate water dikinase, LDH: lactate dehydrogenase, (20) phosphoenolpyruvate carboxykinase (ATP), (21) citrate synthase, (22) aconitate hydratase, IDH: isocitrate dehydrogenase, OFOR: 2-oxoglutarate ferredoxin oxidoreductase, (25) succinyl-CoA synthetase, (26) succinate dehydrogenase, (27) fumarate hydratase, (28) malate dehydrogenase, ME: malic enzyme (oxaloacetate-decarboxylating), (30) aspartate transaminase, (31) asparagine synthetase, (32) L-aspartate oxidase, (33) glutamate synthase, (34) glutamine synthetase, PFOR: pyruvate:ferredoxin oxidoreductase or NifJ: pyruvate-ferredoxin/flavodoxin oxidoreductase, AFOR: aldehyde:ferredoxin oxidoreductase, ADH: alcohol dehydrogenase, (38) phosphate acetyltransferase, (39) acetate kinase.

**Figure 3:**
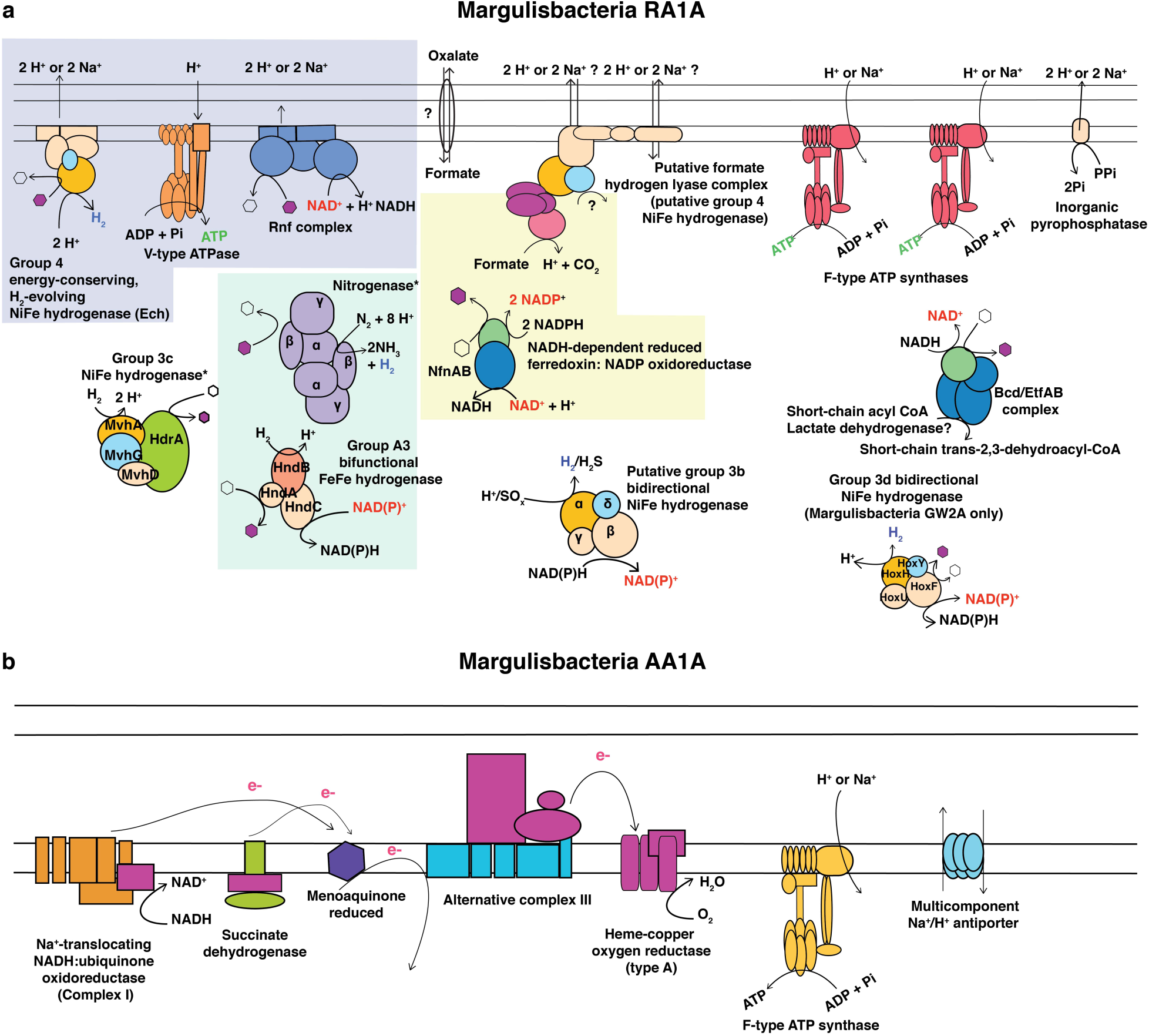
Schematic view of the key protein complexes characteristic of (a) Margulisbacteria RA1A, (b) Margulisbacteria AA1A. Protein complexes encoded by genes in close proximity in the RA1A genome are highlighted as follows: the group 4e NiFe hydrogenase, the V-type ATPase, and the Rnf complex in puple; the nitrogenase and the putative group A3 FeFe hydrogenase in blue; and the putative FHL and Nfn complexes in yellow.

Phylogenetic analyses of the hydrogenase genes suggested that they were horizontally transferred (Figs. 4a and 4b), and that NiFe hydrogenase from groups 3 and 4, considered to be ancestral (Boyd et al., 2014), are encoded in the genome. The great variety of hydrogenases in Riflemargulisbacteria may also reflect adaptation to conditions in anaerobic aquifer environments and allow these bacteria to fine-tune their metabolisms as conditions shift.

**Figure 4a:**
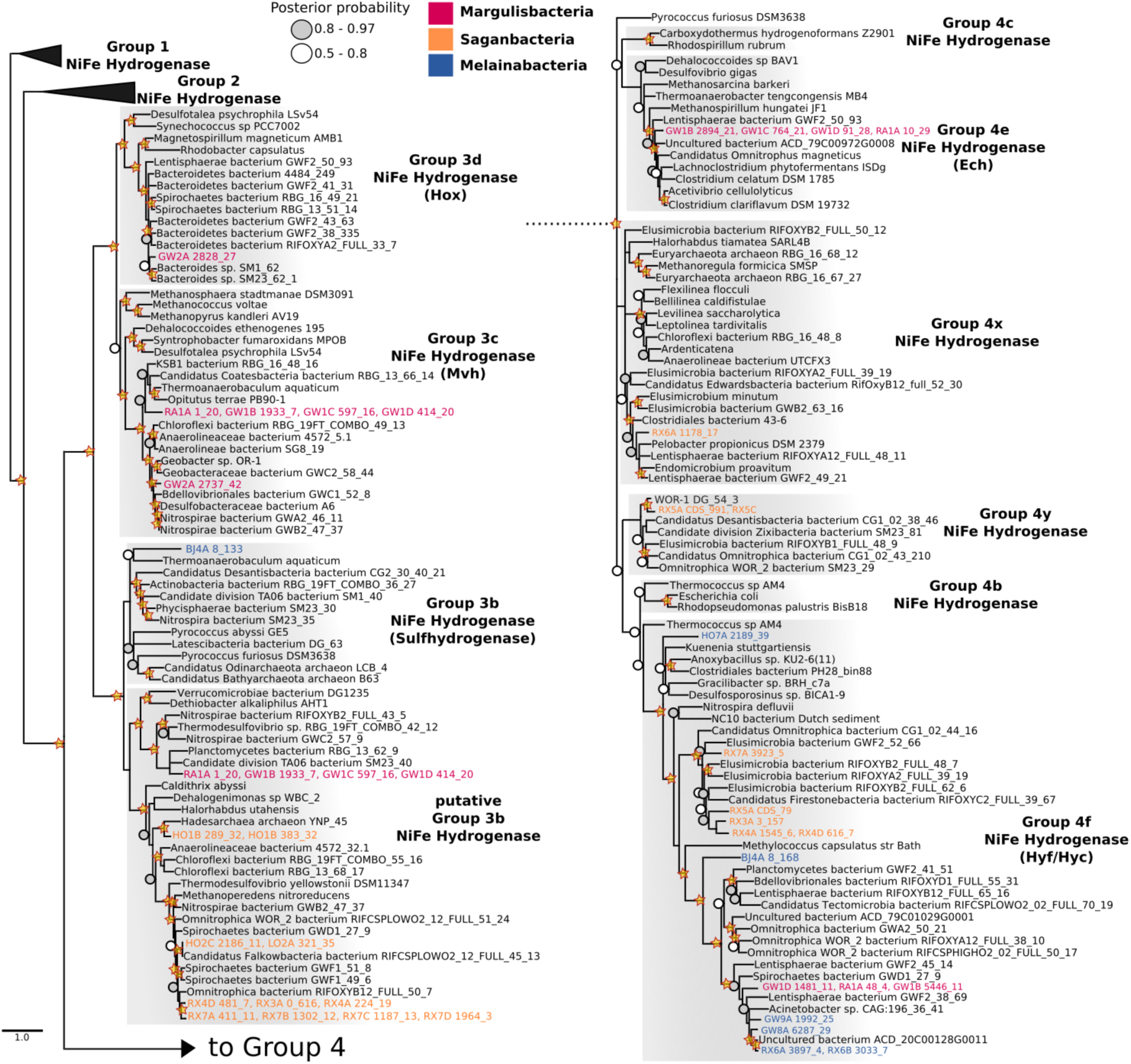
Phylogenetic tree of the catalytic subunit in NiFe hydrogenases. Bayesian phylogeny indicating the phylogenetic relationships of the Riflemargulisbacteria, Saganbacteria, and Melainabacteria NiFe hydrogenase catalytic subunits. Stars highlight branches which were also highly supported (bootstrap support > 97) in a corresponding maximum-likelihood tree (Supplementary Data 1). Tree is rooted at group 1 NiFe hydrogenases. Scale bar indicates substitutions per site. Branches with a posterior support of below 0.5 were collapsed.

**Figure 4b:**
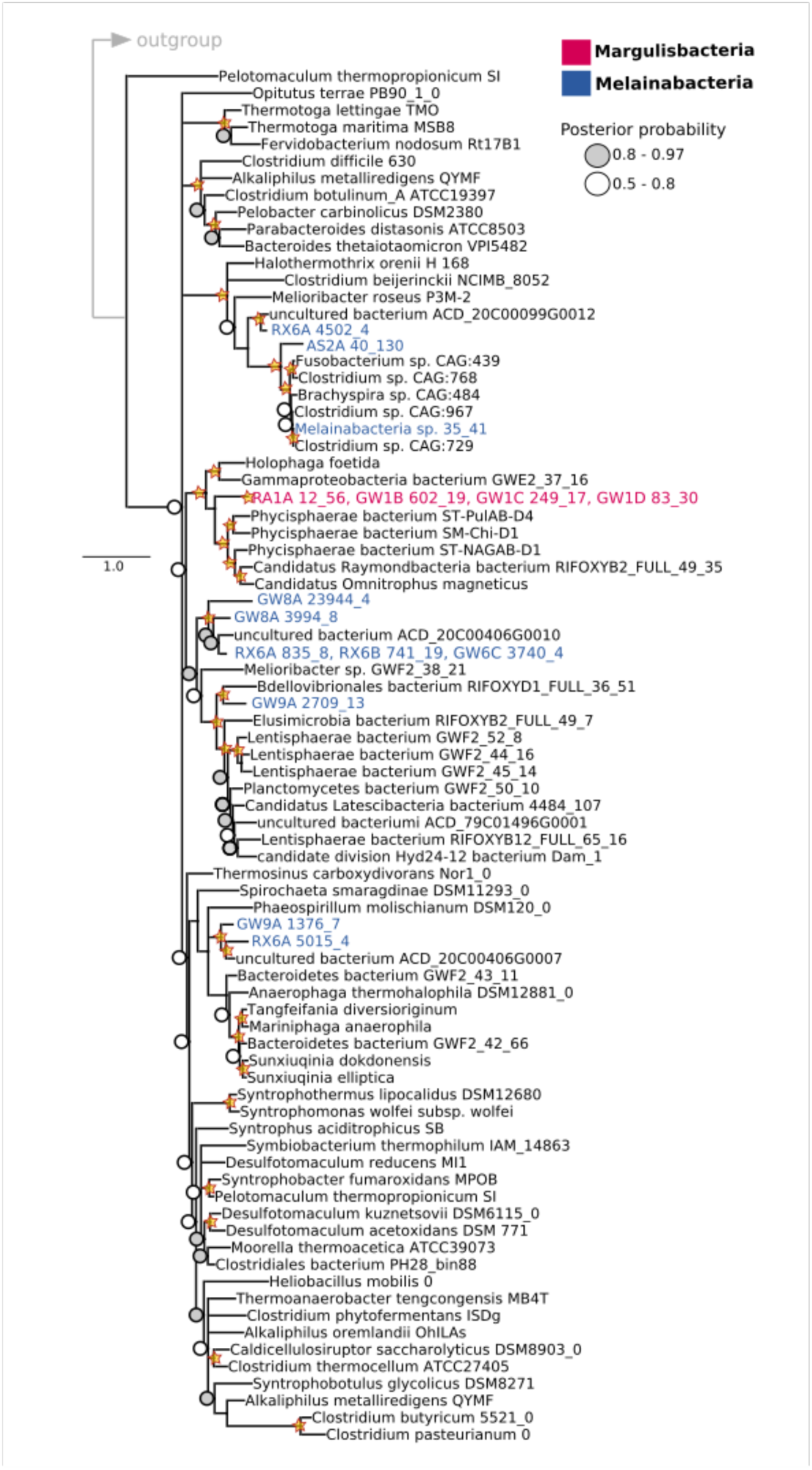
Phylogenetic tree of the catalytic subunit in FeFe hydrogenases. Bayesian phylogeny indicating positions of the Margulisbacteria and Melainabacteria FeFe hydrogenase catalytic subunits. Stars highlight branches which were also highly supported in a corresponding maximum-likelihood tree (Supplementary Data 2). Scale bar indicates substitutions per site. Branches with a posterior support of below 0.5 were collapsed.

We predict that Margulisbacteria RA1A is an obligate anaerobe, as multiple enzymes that participate in central metabolism use ferredoxin as the preferred electron donor/acceptor (Fig. 2). Furthermore, Margulisbacteria RA1A lacks most components of the tricarboxylic acid (TCA) cycle and an ETC (Supplementary Table 2). Carbon dioxide (CO_2_) fixation pathways were not identified in genomes of this lineage (Supplementary Table 2). For further details about the central metabolism of Riflemargulisbacteria see Supplementary Information (Supplementary Text).

Hydrogen metabolism appears essential for the sediment-associated Riflemargulisbacteria. In the absence of a well-defined respiratory electron transport chain, reduced ferredoxin can be re-oxidized by reducing H^+^ to H_2_. Hydrogenases, the key enzymes in hydrogen metabolism, can be reversible and use H_2_ as a source of reducing power, instead of H^+^ as oxidants to dispose of excess reducing equivalents. The assignments of hydrogenase types in this organism were based on phylogenetic analysis of the hydrogenase catalytic subunits (Figs. 4a and 4b) and analysis of the subunit composition.

We identified three types of cytoplasmic NiFe hydrogenases (groups 3b, 3c, and 3d) and one type of cytoplasmic FeFe hydrogenase, in addition to two types of membrane-bound NiFe hydrogenases (groups 4e and 4f) (Fig. 2a). Group 4 hydrogenases in particular are among the most primitive respiratory complexes (Boyd et al., 2014). Membrane-bound group 4 NiFe hydrogenases share a common ancestor with NADH dehydrogenase, a protein complex that couples NADH oxidation with H^+^ or ion translocation (Marreiros et al., 2013). This complex is a key component of respiratory and photosynthetic electron transport chains in other organisms (Friedrich and Weiss, 1997; Friedrich and Scheide, 2000). Given the importance of membrane-bound protein complexes in the generation of a membrane potential in prokaryotic cells, we focused our attention on the membrane-bound hydrogenases found in Riflemargulisbacteria. For a detailed description of cytoplasmic hydrogenases (see Supplementary Text).

The Riflemargulisbacteria are predicted to have two types of membrane-bound H_2_-evolving, ion-translocating group 4 NiFe hydrogenases. One is a group 4e membrane-bound NiFe hydrogenase (Fig. 4a), also called energy-conveting hydrogenases (Ech). These enzymes usually couple the oxidation of reduced ferredoxin to H_2_ evolution under anaerobic conditions (Greening et al., 2016), can be reversible (Meuer et al., 2002), and are typically found in methanogens where they function as primary proton pumps (Welte et al., 2010). Genes for two other complexes are found in close proximity to the Ech-type hydrogenase, a V-type ATPase and a *Rhodobacter n*itrogen *f*ixation (Rnf) electron transport complex (Fig. 2a). These protein complexes may contribute to the generation of a proton motive force in Margulisbacteria RA1A. This is remarkable because both the Ech hydrogenase and the Rnf complex are capable of generating a transmembrane potential, and only a small number of bacteria have been observed to have both (Hackmann and Firkins, 2015). The putative V-type ATPase operon in Margulisbacteria RA1A (*atpEXABDIK*) resembles those of Spirochaetes and Chlamydiales (Lolkema et al., 2003) (Supplementary Table 3). This complex could hydrolyze ATP with H^+^ or Na^+^ translocation out of the cell. Likewise, the Rnf complex, which generally functions as a ferredoxin:NAD^+^ oxidoreductase in *Rhodobacter capsulatum*, couples an exergonic redox reaction to Na^+^ translocation across the membrane (Biegel and Muller, 2010). The Rnf operon in Margulisbacteria RA1A is composed of six genes (*rnfCDGEAB*), similar in organization to the operon in *Acetobacterium woodii* and evolutionarily related to the Na^+^-translocating NADH:quinone oxidoreductase (Nqr) (Biegel et al., 2009; Biegel and Muller, 2010).

Riflemargulisbacteria genomes also encode putative group 4f NiFe hydrogenases, which are energy-converting hydrogenase-related complexes (Ehr). They are putative because they lack the conserved amino acids that coordinate Ni and Fe (Vignais and Billoud, 2007; Marreiros et al., 2013), and function as ferredoxin:NADP^+^ oxidoreductase complexes in *Pyrococcus furiosus* (Mbx complexes) (Schut et al., 2007). Ehr complexes are suspected to couple catalytic reactions in the cytoplasm to charge translocation (Marreiros et al., 2013). In this case, we hypothesize that the group 4f NiFe hydrogenases (Supplementary Fig. 3a) may be part of a formate hydrogenlyase complex that transforms formate into CO_2_ + H_2_ coupled to H^+^/Na^+^ translocation across the membrane. Besides the hydrogenase, additional evidence for the presence of this complex includes a gene for the catalytic subunit (FdhA) of a molybdenum-dependent formate dehydrogenase (Fig. 5) and next to it, two genes encoding FeS cluster-containing subunits. These two genes were annonated as the NADH dehydrogenase electron transfer subunits NuoE and NuoF (Supplementary Fig. 3b; see Supplementary Text).

**Figure 5:**
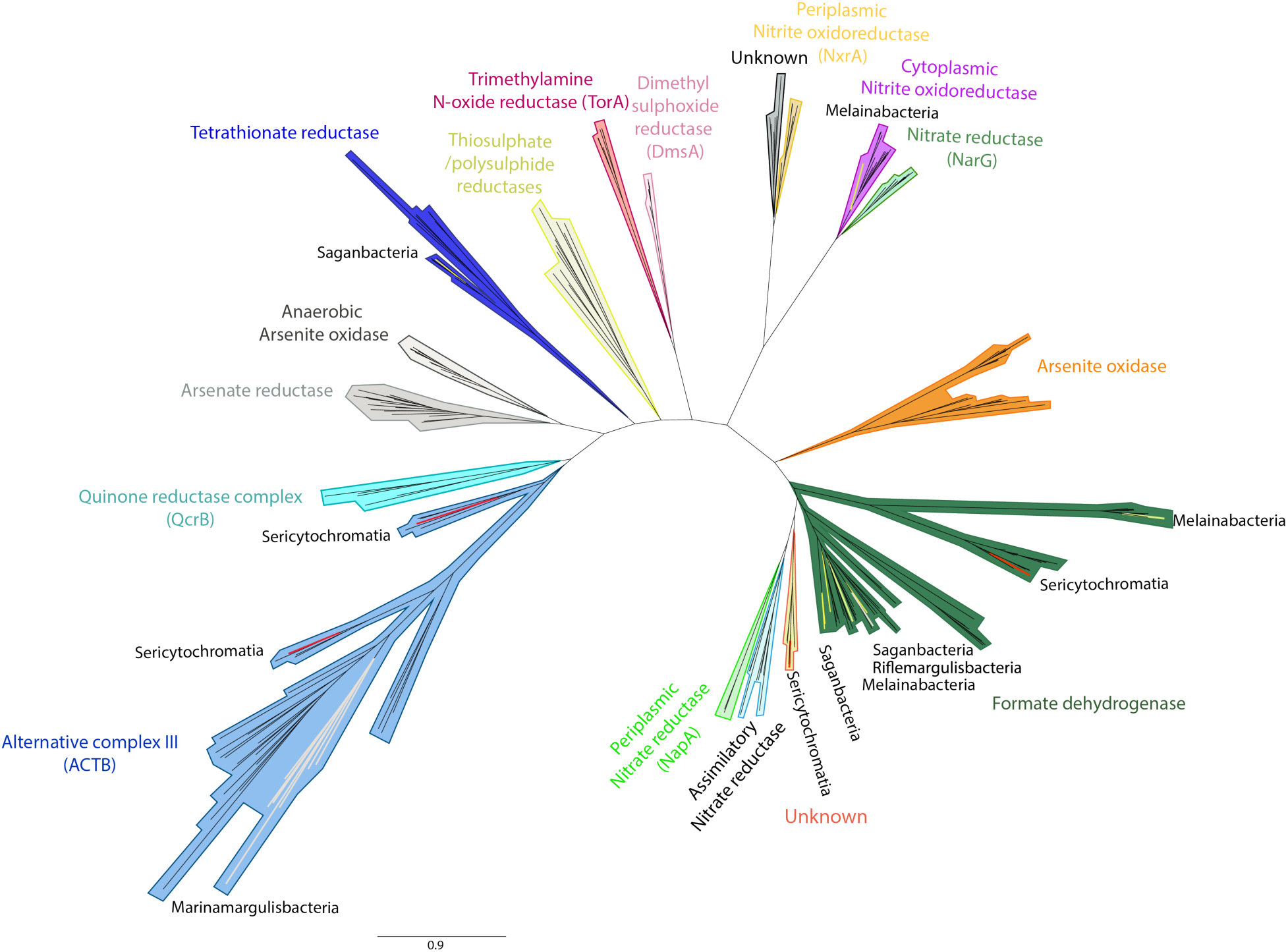
Phylogenetic analysis of the catalytic subunits of the dimethyl sulfoxide (DMSO) reductase superfamily. Catalytic subunits of a formate dehydrogenase (FdhA) were found in Riflemargulisbacteria, Saganbacteria, and Melainabacteria (also present in some Oxyphotobacteria). An Alternative Complex III (ACIII; ActB subunit) was found in Marinamargulisbacteria, as well as in Sericytochromatia LSPB_72 and CBMW_12. Subunit A (TtrA) of a tetrathionate reductase was identified in Saganbacteria. A cytoplasmic nitrate/nitrite oxidoreductase (NXR) was identified in Melainabacteria in this study. The tree is available with full bootstrap values in Newick format in Supplementary Data 3.

Lastly, Margulisbacteria RA1A encodes a nitrogenase, an enzyme known to produce H_2_ during nitrogen fixation. A phylogenetic analysis of NifH (Supplementary Fig. 4) indicates that this enzyme is different from the nitrogenase in Cyanobacteria. Furthermore, we identified genes that encode a bifurcating FeFe hydrogenase (Fig. 4b; putative trimeric group A3, Supplementary Fig. 5) downstream of the *nif* operon. In the forward direction, FeFe hydrogenases can bifurcate electrons from H_2_ to ferredoxin and nincotinamide adenine dinucleotide phosphate –NAD(P)^+^ (Greening et al., 2016). Given the proximity of the FeFe hydrogenase genes on the Margulisbacteria RA1A genome to the Nif operon, this enzyme may be responsible for dissipating excess H_2_ generated by the nitrogenase complex involved in N_2_ fixation.

### Marinamargulisbacteria: metabolic potential for aerobic respiration

Margulisbacteria AA1A, the selected representative genome for the oceanic Marinamargulisbacteria (82% completeness, Supplementary Table 1) shows signs of niche adaptation, including the ability to use O_2_ as a terminal electron acceptor and the lack of hydrogenases. This organism relies on aerobic respiration for generation of a H^+^ potential, and instead of pyruvate fermentation to short-chain fatty acids or alcohols, it may use acetate as a source of acetyl-CoA (Supplementary Table 2a). Phylogenetic analysis of the heme-copper oxygen reductase (complex IV) indicated that the potential for high-energy metabolism was an independently acquired trait (*via* HGT) in this organism (Fig. 6, Supplementary Fig. 6).

**Figure 6:**
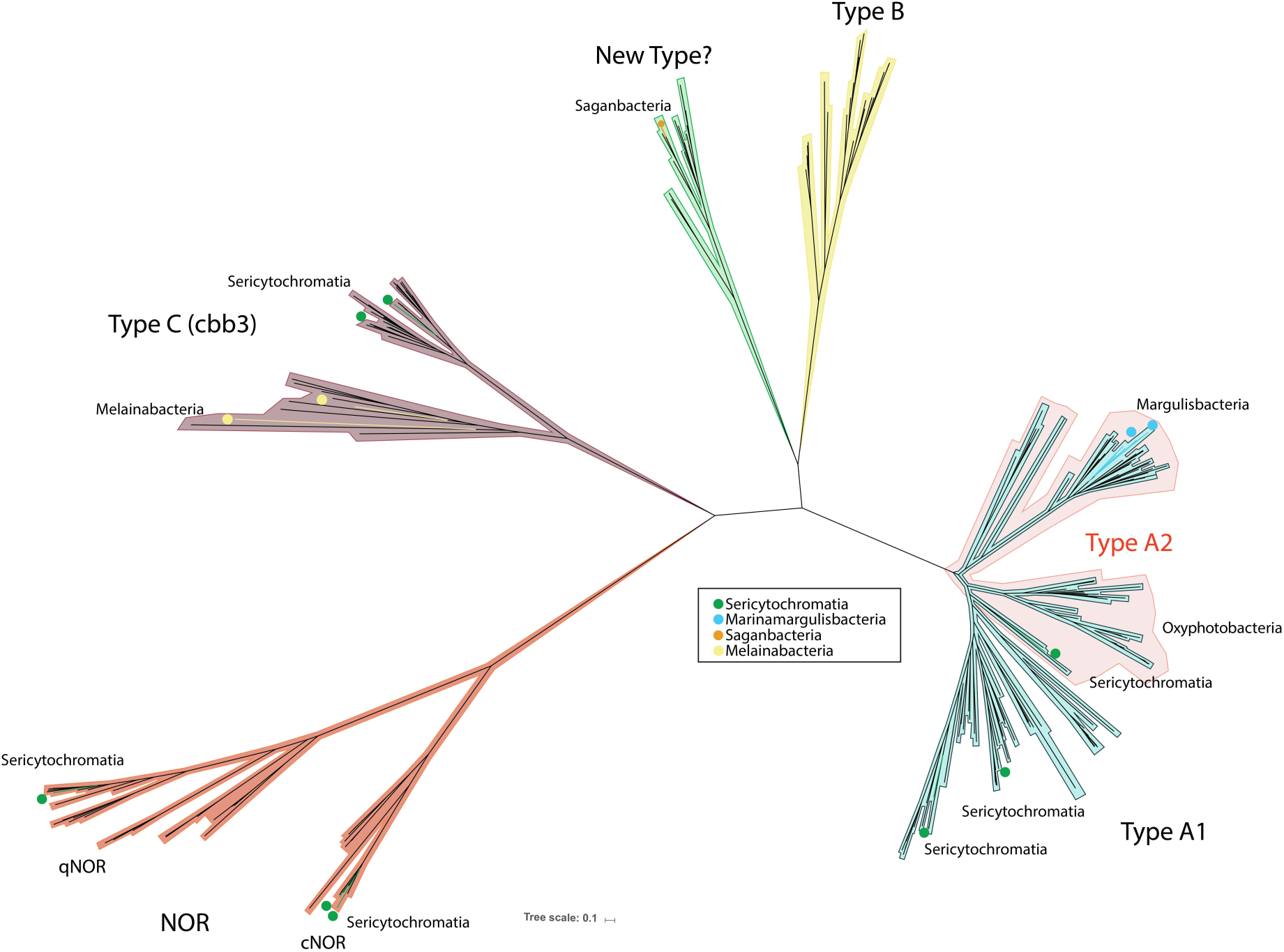
RAxML phylogenetic tree of the catalytic subunit I of heme-copper oxygen reductases. The heme-copper oxygen reductase found in Saganbacteria seems to be a novel type based on phylogenetic analysis of catalytic subunits (CoxA). Marinamargulisbacteria have an A2 type heme-copper oxygen reductase, which is usually found in Oxyphotobacteria. The enzyme in Marinamargulisbacteria is part of a separate phylogenetic cluster from the aa_3_-type heme-copper oxygen reductase found in Oxyphotobacteria (Supplementary Fig. 5). The heme-copper oxygen reductase found in Sericytochromatia LSPB_72 is type A1, and CBMW_12 potentially has two enzymes: one of type A1 and the other of type A2. Sericytochromatia LSPB_72 and CBMW_12 also have type C heme-copper oxygen reductases (CcoN), like Melainabacteria in this study. Two types of bacterial nitric oxide reductase (NOR) have been identified in Sericytochromatia. One is a cytochrome *bc*-type complex (cNOR) that receives electrons from soluble redox protein donors, whereas the other type (qNOR) lacks the cytochrome *c* component and uses quinol as the electron donor. The tree is available with full bootstrap values in Newick format in Supplementary Data 4.

Like the sediment-associated Margulisbacteria, the oceanic Marinamargulisbacteria lack CO_2_ fixation pathways. Thus, we anticipate that they adopt a heterotrophic lifestyle. The genome encodes all enzymes in the TCA cycle (including alpha-ketoglutarate dehydrogenase). NADH produced *via* glycolysis and the TCA cycle must be reoxidized. Unlike Riflemargulisbacteria that have membrane-bound NiFe hydrogenases, in Marinamargulisbacteria AA1A we identified genes encoding a six subunit Na^+^-translocating NADH:ubiquinone oxidoreductase (Nqr; EC 1.6.5.8; *nqrABCDEF*; Supplementary Table 2a). An electron transport chain in Marinamargulisbacteria AA1A (Fig. 2b) could involve electron transfer from NADH to the Nqr, from the Nqr to a menaquinone, then to a quinol:electron acceptor oxidoreductase (alternative Complex III; Fig. 5), and finally to a terminal oxidase (Fig. 6).

Phylogenetic analysis of the gene encoding the catalytic subunit (CoxA) of the terminal oxidase confirms that it is a type A heme-copper oxygen reductase (CoxABCD, EC 1.9.3.1; cytochrome *c* oxidase, (Fig. 6) adapted to atmospheric O_2_ levels. The genes in these organisms are divergent and were probably acquired by horizontal gene transfer based on phylogenetic analysis (Supplementary Fig. 6). The recently released metagenome assembled Marinamargulisbacteria genome UBA6595 from Parks et al. (2017) also encodes an alternative complex III and an type A heme-copper oxygen reductase. In contrast, the Riflemargulisbacteria do not have components of an ETC, except for a single gene of the cytochrome oxidase (*coxB*) presently with unknown function by itself (Supplementary Table 4). Interestingly, we did not identify any hydrogenases in the oceanic Marinamargulisbacteria, probably due to their different habitat.

### Hydrogen metabolism and electron transport chain configurations in Saganbacteria

Like Margulisbacteria, Saganbacteria has anaerobic and aerobic representatives. Similar to sediment-associated Margulisbacteria, Saganbacteria are heterotrophic anaerobes that may generate a H^+^/Na^+^ potential *via* membrane-bound group 4 NiFe hydrogenases, in combination with an Rnf complex. As occurs in the oceanic Margulisbacteria, aerobic organisms within this lineage also rely on an ETC to generate a H^+^ potential. Notably, based on phylogenetic analysis these aerobic organisms encode what looks like a novel type of heme-copper oxygen reductase (Fig. 6), and one organism (HO1A) also encodes the potential for anaerobic respiration (tetrathionate reductase; Fig. 5).

The studied genomes of Saganbacteria encode distinct types of hydrogenases. In the category of cytoplasmic complexes, we only identified one type, the group 3b NiFe sulfhydrogenases, and these occurred in multiple representative genomes (Fig. 4a, Supplementary Table 4). Many Saganbacteria representative genomes also encode three distinct types of membrane-bound hydrogenases (Fig. 4a). Genomic regions including *fdhA* genes also occur in many Saganbacteria with group 4f NiFe hydrogenases (Supplementary Figs. 3b and c), an observation that strengthens the deduction that FHL may occur in these clades.

Another type of group 4 NiFe hydrogenase found in Saganbacteria RX5A is enigmatic.

The most closely related sequences are for hydrogenases found in candidate phyla such as Zixibacteria and Omnitrophica (Fig. 4a). The genomic region encodes a molybdopterin-containing oxidoreductase that could not be classified by phylogeny. Specifically, it could not be identified as a formate dehydrogenase, although the HMMs suggested this to be the case. Also present in the genomic region are a putative CO dehydrogenase subunit F gene (*cooF*), and putative anaerobic sulfite reductase subunits A (*asrA*) and B (*asrB*) (Supplementary Fig. 3e) genes. Overall, the function of the proteins encoded in this genomic region could not be determined (Supplementary Text).

Two other unclassified putative group 4 NiFe hydrogenases were found in Saganbacteria RX6A (Supplementary Figs. 3d and f). Each shares some components with the enigmatic group 4 hydrogenase described above. One of them has antiporter-like subunits and it is phylogenetically most closely related to hydrogenases in *Pelobacter propionicus* and Clostridia (Fig. 4a, Supplementary Fig. 3f).

In terms of their electron transport chain, Saganbacteria HO1A and LO2A possess genes encoding a partial complex I that lacks the NADH binding subunits (NuoEFG) and could use ferredoxin as the electron donor instead (Supplementary Table 4). They also encode a succinate dehydrogenase (SDH) complex that is composed of four subunits and presumably fully functional as a complex II (Supplementary Table 2a). Genes encoding a putative quinol:electron acceptor oxidoreductase complex, which has been previously seen in candidate phyla Zixibacteria (Castelle et al., 2013), may transfer electrons to a terminal reductase and act as a complex III. Quinones, the usual electron donors for this and other respiratory enzymes, were not identified in this genome and alternative biosynthetic pathways should be investigated (Supplementary Table 2a). Remarkably, these genomes also encode for subunits of a quinol oxidase heme-copper oxygen reductase that seems to be a novel type, and that is closely related to a heme-copper oxygen reductase in candidatus *Methylomirabilis oxyfera* (NC10 phylum) (Fig. 6) of unknown type. It is unlikely that these organisms can reoxidize NADH via complex I, and it remains uncertain whether they can use O_2_ as an electron acceptor. Downstream from the genes encoding the putative quinol reductase, there are genes encoding a tetrathionate reductase (Fig. 5), indicating that tetrathionate could be used as a terminal electron acceptor during anaerobic respiration, generating thiosulfate (Hensel et al., 1999). The majority of anaerobic Saganbacteria also possess genes encoding a Rnf complex and a V/A-type ATPase. The V/A-type ATPases have a subunit composition different from that in Riflemargulisbacteria, and their genes are in operons resembling those found in other bacteria and archaea (Supplementary Table 3).

### Hydrogen metabolism and electron transport chain configurations in Melainabacteria

Melainabacteria include organisms capable of fermentation, respiration, or both (Di Rienzi et al., 2013; Soo et al., 2014). In the anaerobic fermentative clades of Melainabacteria, H_2_ plays a central role, and FeFe hydrogenases are more prominent than in Riflemargulisbacteria and Saganbacteria. FeFe hydrogenases generally are more active than NiFe hydrogenases, and are mostly involved in the oxidation of reducing equivalents from fermentation (Bothe et al., 2010). In other clades of Melainabacteria an electron transport chain and a variety of respiratory enzymes can be found, including those for aerobic and anaerobic respiration.

Given the importance of hydrogenases and overall similarity in metabolic capacities of Riflemargulisbacteria and Saganbacteria, we investigated newly available genomes for the Melainabacteria for evidence of yet unrecognized H_2_ metabolism in this group. We found that some representative Melainabacteria genomes have cytoplasmic FeFe hydrogenases that are monophyletic with the FeFe hydrogenase in Margulisbacteria, albeit in a separate phylogenetic cluster (Fig. 4b). These new genomes also encode membrane-bound hydrogenases that could be involved in H_2_ production (Fig. 7). For more detail about cytoplasmic hydrogenases, see Supplementary Text.

**Figure 7:**
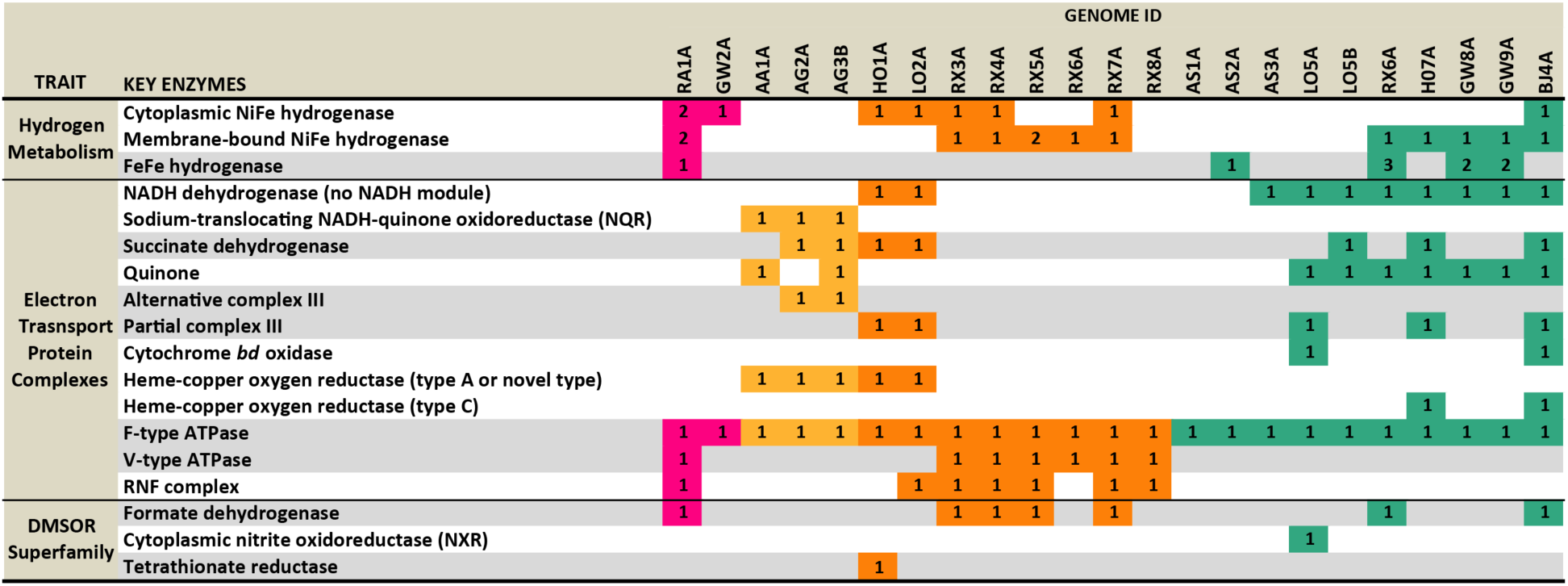
Summary of enzymes involved in hydrogen and energy metabolism in Margulisbacteria, Saganbacteria, and Melainabacteria in this study.

Membrane-bound group 4f NiFe hydrogenases are present in five representative Melainabacteria genomes (Fig. 4a). The sequences in three of the five representative genomes cluster with sequences from Lentisphaerae, Spirochaetes and *Acinetobacter* sp., but Melainabacteria BJ4A branches on its own (Fig. 4a). Melainabacteria HO7A has an entirely different kind of group 4f hydrogenase that is closely related to those of a *Gracilibacteria* sp. and other anaerobic organisms. In addition to membrane-bound group 4f NiFe hydrogenases, Melainabacteria RX6A and BJ4A have an *fdhA* gene, and RX6A has genes encoding NuoE- and NuoF-like subunits just downstream of the *fdhA* gene. Interestingly, FdhA clusters with proteins found in the Cyanobacteria *Nostoc piscinale* and *Scytonema hofmannii* (Fig. 6).

Melainabacteria in this study (except for AS1A and AS1B) have a partial complex I, similar to the one in Saganbacteria (Fig. 7, Supplementary Table 4). Melainabacteria, represented by genomes LO5B, HO7A and BJ4A, encode a partial SDH (complex II).

Intriguingly, cytochrome *b*_*6*_ (PetB; KEGG ortholog group K02635) and the Rieske FeS subunit (PetC; K02636) of the cytochrome *b*_*6*_*f* complex (complex III) were identified by HMMs predictions in these genomes. Only Melainabacteria LO5B, HO7A, and BJ4A are potentially able to use O_2_ and other terminal electron acceptors. Specifically, Melainabacteria HO7A and BJ4A harbor O_2_ reductases (see Supplementary Text) in the vicinity of genes encoding part of the cytochrome *b*_*6*_*f* complex (*petB* and *petC*), suggesting that they have a complex III/IV combination. Whether Melainabacteria are able to synthesize ubiquinone remains to be determined because only five out nine genes required for its synthesis were identified in the genomes of sediment-associated Melainabacteria (Supplementary Table 2a).

Other forms of respiration were predicted in some Melainabacteria in this study. For instance, Melainabacteria LO5A encodes a cytoplasmic nitrite/nitrate oxidoreductase (Fig. 5). Nitrite/nitrate oxidoreductases (NXR) participate in the second step of nitrification, the conversion of nitrite to nitrate and can also be reversible (Lücker et al., 2010). These genes are followed by another gene encoding cytochrome *b*_*6*_ (*petB*) and a nitrate/nitrite transporter (*narK*; K02575).

### Additional complexes possibly involved in electron bifurcation

Some Margulisbacteria RA1A, Saganbacteria and Melainabacteria have complexes predicted to be involved in electron bifurcation. Electron bifurcation is the most recently described mechanism for energy conservation (the alternative mechanisms are substrate-level phosphorylation and electron transport phosphorylation) (Buckel and Thauer, 2013). For hydrogenase-based electron bifurcation, reduced ferredoxin and NAD(P)H (redox carriers with very different electrode potentials) are oxidized, coupled to the reduction of H^+^ to form H_2_ (Peters et al., 2016). This reaction can also work in the reverse as the electron bifurcation complex is bidirectional. Electron bifurcation complexes allow balance of reduced and oxidized cofactors in anaerobic environments, where electron acceptors are limited (Boyd et al., 2014).

In close proximity to the region encoding FdhA in Margulisbacteria RA1A and most Saganbacteria genomes, we identified genes encoding a NADH-dependent reduced ferredoxin: NADP oxidoreductase (Nfn) complex (Fig. 2a). Nfn complexes use 2 NADPH molecules to catalyze the reversible reduction of oxidized ferredoxin and NAD^+^ (Demmer et al., 2015), thus it is involved in electron bifurcation.

Margulisbacteria RA1A, all Saganbacteria and many Melainabacteria have genes encoding a putative butyryl-CoA dehydrogenase (Bcd) in close proximity to genes encoding electron transfer flavoprotein subunits EtfA and EtfB (Fig. 2a). Together, these may form a Bcd/EtfAb complex, which is usually involved in electron bifurcation reactions between crotonyl-CoA, ferredoxin and NADH. Specifically, Bcd catalyzes the transformation of short-chain acyl-CoA compounds to short-chain trans-2,3-dehydroacyl-CoA using electron-transfer flavoproteins as the electron acceptor (EtfAB). Alternatively, this complex in Margulisbacteria RA1A could be coupled to oxidation through lactate dehydrogenase under conditions of low carbon availability (see Supplementary Text).

## Discussion

Contemporary non-photosynthetic relatives of the Oxyphotobacteria share several interesting genomic features that frame photosynthesis in the context of metabolisms that likely existed prior to its evolution. These include numerous hydrogenases, widespread lack of an oxidative phosphorylation pathway, and lack of carbon dioxide fixation pathways. Against this background, photosynthesis represents a dramatic net-energy gain and access to a different life strategy – one that does not rely on limited organic carbon as a source of electrons or complexes that provide only low-energy yields linked to ion translocation.

We set out to study the metabolic potential of the phylogenetic relatives of Oxyphotobacteria and identified genomic traits that may have been present in their common ancestor. The genomes described here present a more detailed picture than was previously available of the metabolic platform in which oxygenic photosynthesis and aerobic respiration evolved. Based on our analyses, the common ancestor of Margulisbacteria, Saganbacteria, Melainabacteria, Sericytochromatia, and Oxyphotobacteria was likely an anaerobe with fermentation-based metabolism. Even today, Oxyphotobacteria can switch to fermentative metabolism under anoxic conditions, and at the heart of this process is a bidirectional group 3d NiFe hydrogenase (Appel, 2012).

### Hydrogen metabolism may be a common thread in the evolutionary history of Oxyphotobacteria and sibling lineages

Anaerobic representatives of Riflemargulisbacteria and Saganbacteria have multiple protein complexes that participate in electron-transport phosphorylation (ETP) and electron bifurcation (EB) in the context of H_2_ metabolism. In Riflemargulisbacteria and Saganbacteria, membrane-bound group 4 NiFe hydrogenases may have a respiratory function. These protein complexes could make up a minimal, highly efficient respiratory chain, as previously suggested for these types of enzymes (Greening et al., 2016). Riflemargulisbacteria is unique among the anaerobic lineages because its genome is predicted to encode multiple cytoplasmic H_2_-consuming hydrogenases, in addition to membrane-bound energy-conserving H_2_-evolving NiFe hydrogenase (Ech; Group 4e), an Rnf complex, and potentially a FHL complex. The cytoplasmic hydrogenases constitute the main mechanism by which these bacteria oxidize reducing equivalents that result from core metabolic pathways, and produce H_2_. The main function of the Ech hydrogenase and the Rnf complex is to couple ferredoxin oxidation with ion-translocation; and a V-type ATPase could be involved in ATP synthesis from the ion potential generated from both these complexes. Anaerobic Saganbacteria that also possess an Rnf complex and a V-type ATPase may be able to use this mechanism for energy conservation as well. Furthermore, two groups of closely related Saganbacteria have group 4 hydrogenases with unknown electron donors that may also be involved in the creation of a H^+^/Na^+^ potential. Thus, Saganbacteria could still produce energy *via* group 4 NiFe hydrogenases.

The the putative formate hydrogenlyase (FHL) complex in anaerobic Riflemargulisbacteria and Saganbacteria is highly unusual. We speculate that this putative FHL may be an ancient form of the complex and involved in oxidative phosphorylation. Under high extracellular formate concentrations, the *E. coli* FHL complexes release electrons from formate oxidation that are passed down other peripheral subunits containing FeS-binding clusters to the site for H_2_ formation (Pinske and Sawers, 2016). Of the two different hydrogenases that are considered to be part of an FHL complex in *E. coli*, only hydrogenase 3 (encoded by *hyc* genes) has been purified (McDowall et al., 2014). Moreover, ion-translocation has only been observed in similar complexes in the genus *Thermococcus* (Kim et al., 2010). However, we suspect that in Riflemargulisbacteria and Saganbacteria the Nuo-like electron transfer subunits together with the small and large NiFe hydrogenase subunits pass electrons down to the transmembrane subunits, where ion translocation takes place. For other Ehr complexes (*e.g.,* Mbx and Mbh hydrogenases), it was suggested that the distance between the formate dehydrogenase and membrane-bound subunits is important for the creation of an electrochemical potential across the membrane (Marreiros et al., 2013). Alternatively, the putative group 4f hydrogenase in these organisms may interact with an unknown substrate that allows for an electron bifurcation mechanism.

Riflemargulisbacteria, Saganbacteria, and Melainabacteria all have putative group 4f membrane-bound NiFe hydrogenases that lack the known catalytic residues, possibly due to the divergence between these sequences and hydrogenases that have been studied experimentally. Experimental testing is needed to determine whether these hydrogeases are part of FHL complexes, are expressed, assembled and functional. The existence of FHL complexes that might include the group 4f hydrogenase is even more questionable in Melainabacteria BJ4A, because the genes encoding the electron transfer subunits were not identified. Such complexes were not identified in previously described Melainabacteria either (Di Rienzi et al., 2013). Regardless of the true function of the putative group 4f membrane-bound NiFe hydrogenases, it is interesting to discover the variety of other membrane-bound energy-conserving complexes in lineages sibling to Oxyphotobacteria. Membrane-bound group 4 NiFe hydrogenases share a common ancestor with the NADH dehydrogenase (Complex I) and the FHL complex, in which the catalytic subunits of NiFe hydrogenases may have later become the electron transfer subunits of complex I (Friedrich and Scheide, 2000; Marreiros et al., 2013).

Notable in Riflemargulisbacteria in particular (but present in other groups) are a variety of potential electron bifurcating complexes, including the Nfn and Bcd/Etf complexes (Lubner et al., 2017). Although not described as such, cytoplasmic NiFe hydrogenases (groups 3c and 3d) and cytoplasmic bidirectional FeFe hydrogenases could be involved in electron bifurcation.

Overall, we identified potentially six electron bifurcation complexes in Riflemargulisbacteria. These are common in fermentative organisms, but the number and variety found in Riflemargulisbacteria is notable. These complexes probably serve as the primary mechanism to balance of redox carriers in these organisms.

### Aerobic and anaerobic electron transport chains, the gain of high-energy metabolism

From the perspective of metabolic evolution, it is interesting to consider the origin of high electrode potential electron transport chains and their components. In this study, we were particularly interested in the electron entry (complex I) and exit points (terminal oxidases).

Within the Saganbacteria and Melainabacteria, Saganbacteria HO1A and LO2A and Melainabacteria LO5B, HO7A and BJ4A are the only organisms that posses some sort of an electron transport chain. Given the prediction that the common ancestor of all groups was a fermentative anaerobe, we suspect that the ETC and ability to use O_2_ was a later acquired trait.

Saganbacteria HO1A and LO2A and most Melainabacteria in this study (except AS1A and AS2A) have in common with Oxyphotobacteria the lack of the NADH module of NADH dehydrogenase (complex I). Respiration in the thylakoid membrane of Oxyphotobacteria is mostly initiated from succinate rather than NADH. However, how succinate dehydrogenase (SDH; complex II) works in *Synechocystis* sp. is not completely understood, in part due to the lack of two (out of four) subunits (Vermaas, 2001; Lea-Smith et al., 2016). This is also the case in Melainabacteria LO5B, HO7A and BJ4A. The complex I found in Saganbacteria HO1A and LO2A and Melainabacteria may be involved in proton translocation independent of NADH oxidation, and thus it may play a role similar to that of membrane-bound NiFe hydrogenases.

Remarkably, Saganbacteria HO1A and LO2A also carry a putative novel type of heme-copper oxygen reductase (quinol oxidase), which could act as the terminal reductase in an electron transport chain.

Only Marinamargulisbacteria is predicted to have an electron transport chain that includes an alternative complex III (ACIII) and a type A heme-copper oxygen reductase (cytochrome oxidase adapted to high O_2_ levels), which may act as the intermediary between complex I and the terminal oxidase. Marinamargulisbacteria UBA6595 (Parks et al., 2017) and Sericytochromatia CBMW_12 (Soo et al., 2017) are the only other related bacteria known to have an ACIII and a type A heme-copper oxygen reductase. Phylogenetic analyses indicate that Marinamargulisbacteria and Sericytochromatia acquired this complex by lateral gene transfer. Given that Marinamargulisbacteria AA1A was found in the ocean, where dissolved O_2_ may be available, it must have evolved the necessary machinery (*i.e.*, complex I, menaquinone, ACIII, cytochrome oxidase) to take advantage of O_2_ as a terminal electron acceptor. Similarly, fermentation and H_2_ metabolism may be less relevant in the water column, and this clade may have lost the ancestral traits that were advantageous in other redox conditions.

The type C heme-copper oxygen reductase (complex IV) found in Melainabacteria HO7A and BJ4A may be adapted to low O_2_ levels, as expected for microaerophilic or anoxic environments such as the subsurface and the human gut. Some aerobic Melainabacteria studied here, as well as *Obscuribacter phosphatis* (Soo et al., 2014; Soo et al., 2017), have a fusion involving complex III and complex IV. However, we annotated the proteins related to complex III in these Melainabacteria as cytochrome *b*_*6*_*f* subunits. If correct, this is important because complex *b*_*6*_*f* has only been found to play the role of complex III in Oxyphotobacteria (Soo et al., 2017). Thus, cytochrome *b*_*6*_*f* complex may have been present in the common ancestor of all of these lineages.

Notably, Melainabacteria BJ4A could have a branched electron transport chain, with one branch leading to a cytochrome *d* ubiquinol oxidoreductase and the other leading to the type C heme-copper oxygen reductase. Both these enzymes have high affinity for O_2_, thus are adapted to low O_2_ levels. This is unusual considering that when two O_2_ reductases are present in other lineages studied here and Oxyphotobacteria they have different O_2_ affinities. For instance, Oxyphotobacteria have a type A heme-copper oxygen reductase (low O_2_ affinity, adapted to high O_2_ levels) in the thylakoid membrane and a quinol cytochrome *bd* oxidase (high affinity for O_2_) in the cytoplasmic membrane (in addition to a cytochrome *bo* oxidase) (Vermaas, 2001).

Sericytochromatia also have two types of heme-copper oxygen reductases, type A and type C, that also differ in their affinity for O_2_ (low and high affinity, respectively).

### Absence of genes involved in carbon fixation and photosynthesis in linages sibling to Oxyphotobacteria

Based on the absence of genes for CO2 fixation and photosynthetic machinery in Melainabacteria and Sericytochromatia, the lineages most closely related to Oxyphotobacteria, it was suggested that these capacities arose after their divergence (Di Rienzi et al., 2013; Shih et al., 2017; Soo et al., 2017). However, the alternative possibility is that they were a characteristic of their common ancestor, but lost in Melainabacteria and Sericytochromatia. The current analyses support the former conclusion, given the essentially complete lack of genes that may be involved in carbon fixation and photosynthesis in two additional lineages sibling to Melainabacteria, Sericytochromatia, and Oxyphotobacteria.

### Concluding remarks

There is considerable interest in the metabolism of lineages sibling to the Cyanobacteria.

Genomes from these lineages may provide clues to the origins of complexes that could have evolved to enable aerobic (and other types of) respiration *via* an electron transport chain. Based on the analyses presented here, we suggest that H_2_ was central to the overall metabolism, and hydrogenases and the Rnf complex played central roles in proton translocation for energy generation, with redox carrier balance reliant upon electron bifurcation complexes. Overall, members of the lineages related to Oxyphotobacteria appear to have evolved to optimize their metabolisms depending on their redox environment.

## Methods

### Samples collection, DNA extraction, and sequencing

Genomes in this study were recovered from several sources (Supplementary Table 1). Margulisbacteria GW1B-GW1D, Saganbacteria HO1A, HO1B, LO2A, LO2B, HO2C, RX3A-RX8C, Melainabacteria HO7A, LO5A, LO5B, RX6A-RX6C, GW8A, and GW9A originated from an alluvial aquifer in Rifle, CO, USA (groundwater samples). For sampling, DNA processing, sequencing information, metagenome assembly, genome binning and curation of these aquifer genomes see Brown, et al. (2015) and Anantharaman, et al. (2016).

Margulisbacteria RA1A was obtained from an acetate amendment experiment (Rifle Acetate Amendment Columns) conducted at Rifle in 2010 (Handley et al., 2015). Individual columns packed with sediment (13 and a background sample) were sacrificed from days 13 to 61 to encompass two geochemical processes: iron reduction and sulfate reduction (Kantor et al., 2013). DNA extraction, Illumina library preparation and DNA sequencing protocols were described elsewhere (Wrighton et al., 2012; Handley et al., 2015).

Margulisbacteria AA1A and AG2A-AG6A were collected from the Gulf of Maine at a 1 m depth off the coast of Boothbay Harbor, Maine (43.84 N, −69.64 W) on 16 September, 2009 (AAA071) and sorted immediately, targeting small heterotrophic protists, as described previously (Martinez-Garcia et al., 2012). 1 mL samples were collected at a depth from 112 m and 180 m the Southeast Pacific Ocean (−23.46 N, −88.77 E and −26.25 N, −103.96 E for AG-333 and AG-343, respectively) on 1 December, 2010, from the Western Atlantic Ocean at a depth of 100 m (9.55 N, −50.47 E and 24.71 N, −66.80 E for AG-410 and AG-414, respectively) on 1 June, 2010, and at a depth of 72 m in the Eastern Atlantic (17.40 N, −24.50 E) on 1 December, 2011 (AG-439), amended with 10% glycerol and stored at −80°C.

Melainabacteria BJ4A was obtained from the Mizunami Underground Research Laboratory in Japan. For sampling refer to Ino et al. (2017) and for DNA processing and sequencing methods refer to Hernsdorf et al. (2017). In brief, groundwater (30 L) was collected in 2014 from a highly fractured bedrock domain (HFDB) in a horizontal borehole (interval #3) part of a 300 m deep stage, by filtration through 0.22 μm GVWP filters. Genomic DNA was extracted using an Extrap Soil DNA Kit Plus version 2 (Nippon Steel and Sumikin EcoTech Corporation), libraries prepared using TruSeq Nano DNA Sample Prep Kit (Illumina), and 150 bp PE reads with a 550 bp insert size were sequenced by Hokkaido System Science Co. using Illumina HiSeq2500 (Hernsdorf et al., 2017). Data assembly and genome binning was carried out as described in Hernsdorf et al. (2017).

Four other Melainabacteria genomes (AS1A, AS2A, AS3A, and AS3B) were part of a study of the microbiome in the arsenic-impacted human gut. Faecal samples were obtained from 10 Bangladeshi men (aged between 27-52) from the Laksam Upazila, Bangladesh, that all displayed signs of arsenicosis and were consuming arsenic in their drinking water. Samples were collected on four consecutive days and stored at −20°C until they were shipped to the UK on dry ice. The samples were then stored at −80°C until nucleic acid extraction. DNA was isolated from the faecal samples using PowerFaecal DNA isolation kit (MoBio) according to the manufacturer’s instructions and stored at −20°C until they were sent to RTLGENOMICS (Texas, USA) frozen on dry ice. Samples were then prepared using the Kapa HyperPlus Kit (Kapa Biosystems) following the manufacturer’s protocol, except that the DNA was fragmented physically using the Diagenode Bioruptor, instead of enzymatically. The resulting individual libraries were run on a Fragment Analyzer (Advanced Analytical) to assess the size distributions of the libraries, quantified using a Qubit 2.0 fluorometer (Life Technologies), and also quantified using the Kapa Library Quantification Kit (Kapa Biosystems). Individual libraries were then pooled equimolar into their respective lanes and loaded onto an Illumina HiSeq 2500 (Illumina, Inc.) 2 × 125 flow cell and sequenced.

### Data processing, assembly and binning

*Margulisbacteria RA1A:* Reads for each of the 14 RAAC samples were first trimmed using sickle (https://github.com/najoshi/sickle) and assembled using idba_ud (Peng et al., 2012) with default parameters. To bin the assembled data we used an abundance-pattern based approach in which scaffolds are clustered based on their abundance across multiple samples (Sharon et al., 2013) using ESOM (Ultsch and Mörchen, 2005; Dick et al., 2009). Reads were mapped separately to each of the 14 assemblies using bowtie (Langmead et al., 2009) with parameters -q -n 2 -e 200 -p 6 --best --sam. To prepare the data for ESOM we used version 1.00 of the script prepare_esom_files.pl (https://github.com/CK7/esom/blob/master/prepare_esom_files.pl) with the 14 read mapping files for each assembly and window and minimum size parameters both set to 3000 bps. Binning itself was done manually for each of the 14 ESOM maps. Overall we recovered 361 bins.

Description of the rest of the data will be provided elsewhere. Bin for Margulisbacteria RAAC was recovered from the aac11 sample, during the sulfate reduction phase (Kantor et al., 2013).

*Melainabacteria AS1A-AS3B:* Reads for each of the arsenate gut samples were trimmed and assembled using the same method as Margulisbacteria RAAC (see above). Samples were binned using DASTool v1.0 (Sieber et al.) with input bins generated using CONCOCT v0.4.1

(Alneberg et al., 2014), ABAWACA v1.00 (https://github.com/CK7/abawaca), MaxBin v2.2 (Wu et al., 2016), and the ggKbase binning interface (Raveh-Sadka et al., 2015). For AS1A and AS2A, GC content, coverage, and phylogenetic profile were used for refinement of the genomes.

### Curation of genomes derived from metagenomes

*Margulisbacteria RA1A:* The ESOM bin included 147 scaffolds. The bin was then manually curated by using a mini-assembly approach as described in Sharon et al. (2013). The mini-assembly process is iterative and was done manually as follows. First, the reads from the aac11 sample are mapped to the bin scaffolds using bowtie with parameters as described above. Next, for each scaffold end, all reads that align to the end are collected along with their pair-end mates. These reads are then assembled and the scaffold is elongated based on this local assembly. Elongation process stops once either one or more other scaffolds are linked to the end or when no local assembly can be generated, usually due to low or no coverage. The process is repeated until no further elongation can be done to any of the scaffolds. 23 mini-assembly iterations were performed overall. Resulting number of scaffolds is 111, with a total length of 3.43 mbp. As was discovered during the process, the high number of scaffolds is due to the presence of close to 80 transposase and integrase genes in the genome.

Assembly errors were identified and repaired using a previously described method (https://github.com/christophertbrown/fix_assembly_errors/releases/tag/2.00) (Brown et al., 2015). Briefly, errors were identified as regions of no coverage by stringently mapped paired-read sequences. Stringent mapping was conducted by first mapping reads to the assembly using Bowtie2 (Langmead and Salzberg, 2012) with default parameters for paired-reads. The software repairs assembly errors using Velvet (Zerbino and Birney, 2008), to re-assemble reads that map to those regions based on less-stringent criteria (only one read in the pair has to map to a particular region with less than two mismatches). The final assembly was visually inspected with mapped paired read sequences using Geneious (Kearse et al., 2012). As a result of this process, about 98% of the genes remained unchanged. Contigs whose phylogenetic affiliation did not match that of the majority were removed from the genome.

*Melainabacteria BJ4A:* Local assembly errors were manually identified and repaired and the genome was curated in Geneious. Paired reads were used to full gaps and to extend and join scaffolds.

*Melainabacteria AS1A-AS3B:* Local assembly errors were identified and repaired using the previously described method (Brown et al., 2015).

### Single-cell genomics

The generation, identification, sequencing and assembly of single amplified genomes (SAGs) was performed at the Bigelow Laboratory Single Cell Genomics Center (scgc.bigelow.org). Samples were initially passed through a 70 μm (AAA071) or 40 μm (others) strainer (Becton Dickinson) and incubated for 10–60 min with LysoTracker Green (AAA071) or SYTO-9 (others) stains (Thermo Fisher Scientific). Fluorescence-activated cell sorting (FACS) was performed with the BD MoFlo (AAA071) or InFlux Mariner (others) flow cytometers equipped with 488 nm lasers and 70 μm nozzle orifices (Becton Dickinson). Cytometers were triggered to sort on side scatter, and the “single-1 drop” mode was used for purity. For each sample, individual cells were deposited into 384-well plates containing 600 nL per well of 1x TE buffer and stored at −80°C until further processing: 315 or 317 wells were dedicated for single particles, 66 or 64 wells were used as negative controls (no droplet deposition), and 3 wells received 10 particles each to serve as positive controls. The DNA for each cell was amplified using MDA (AAA071) or WGA-X (others), as previously described in Stepanauskas et al. (2017). 16S RNA genes were analyzed as in in Stepanauskas et al. (2017). Shotgun Illumina libraries were created and sequenced, and the reads were assembled using SPAdes v3.9.0 (Bankevich et al., 2012) with modifications, as previously described (Stepanauskas et al., 2017). Three bacterial benchmark cultures were used to evaluate SAGs for assembly errors. Benchmark cultures have diverse genome complexity and %GC, resulting in 60% average genome recovery, no non-target and undefined bases, and average frequencies of misassemblies, indels and mismatches per 100 kbp: 1.5, 3.0, and 5.0 (Stepanauskas et al., 2017). Bacterial symbiont contigs of the protist SAG AAA071-K20 were identified and separated from the host genome using tetramer principal component analysis, relying on the first two principal components (Woyke et al., 2009). Final SAG assemblies are deposited in the Joint Genome Institute Integrated Microbial Genomic database (https://img.jgi.doe.gov/).

### Completeness estimation

Genome completeness was evaluated based on a set of 51 single copy genes previously used (Anantharaman et al., 2016). Genomes from metagenomes and single-cell genomes were categorized according to the minimum information about a metagenome-assembled genome (MIMAG) and a single amplified genome (MISAG) of bacteria and archaea (Bowers et al., 2017).

### Functional annotations

Predicted genes were annotated using Prodigal (Hyatt et al., 2010), and similarity searches were conducted using BLAST against UniProtKB and UniRef100 (Suzek et al., 2007), and the Kyoto Encyclopedia of Genes and Genomes (KEGG) (Ogata et al., 1999) and uploaded to gg Kbase (http://www.ggkbase.berkeley.edu). Additionally, the gene products were scanned with hmmsearch (Finn et al., 2011) using an *in house* Hidden Markov Models (HMMs) database representative of KEGG orthologous groups. More targeted HMMs were also used to confirm the annotation of genes encoding hydrogenases (Anantharaman et al., 2016).

### Gene content comparison

Inference of clusters of orthologous proteins was performed with OrthoFinder 2.1.3 (Emms and Kelly, 2015) on a set of genomes representing the following class and phylum level lineage-specific pangenomes: Margulisbacteria (n=14), Saganbacteria (n=26), Sericytochromatia (n=3), Melainabacteria (n=17) and Cyanobacteria (n=34). Pangenomes include all proteins found in the respective set of genomes. Cyanobacterial genomes were de-replicated so that for each genus available in IMG (Chen et al., 2016) one genome was included in the dataset. Shared protein families between lineage pangenomes and lineage-specific protein families were identified based on the OrthoFinder orthogroups table. For each pairwise pangenome comparison the average percentage of shared protein families was calculated based on the total number of shared protein families divided by the sum of non-lineage specific protein families found in the two pangenomes. The pangenome comparisons were visualized using the python packages matplotlib and pyUpSet (https://github.com/ImSoErgodic/py-upset).

### Phylogenomics

Two different species trees were constructed, the first tree to place Sagan- and Margulisbacteria into phylogenetic context with other bacterial phyla and the second one to provide a more detailed view of taxonomic placement of different Sagan- and Margulisbacteria. To reduce redundancy in the first species tree DNA directed RNA polymerase beta subunit 160kD (COG0086) was identified in reference proteomes, Saganbacteria and Margulisbacteria using hmmsearch (hmmer 3.1b2, http://hmmer.org/) and the HMM of COG0086 (Tatusov et al., 2000). Protein hits were then extracted and clustered with cd-hit (Fu et al., 2012) at 65% sequence similarity. Cluster-representatives were used to build the species tree of the Terrabacteria. For the detailed species tree (Fig. 1b), pairwise genomic average nucleotide identity (gANI) was calculated using fastANI (Jain et al., 2017). Only genome pairs with an alignment fraction of greater than 70% and ANI of at least 98.5% were taken into account for clustering with mcl (Enright et al., 2002). Representatives of 98.5% similarity ANI clusters were used for tree building based on two different sets of phylogenetic markers; a set of 56 universal single copy marker proteins (Eloe-Fadrosh et al., 2016; Yu et al., 2017) and 16 ribosomal proteins (Hug et al., 2016). For every protein, alignments were built with MAFFT (v7.294b) (Katoh and Standley, 2013) using the local pair option (mafft-linsi) and subsequently trimmed with BMGE using BLOSUM30 (Criscuolo and Gribaldo, 2010). Query genomes lacking a substantial proportion of marker proteins (less than 28 out of 56) or which had additional copies of more than three single-copy markers were removed from the data set. Single protein alignments were then concatenated resulting in an alignment of 13849 sites for the set of 56 universal single copy marker proteins and 2143 sites for the 16 ribosomal proteins. Maximum likelihood phylogenies were inferred with IQ-tree (multicore v1.5.5)(Nguyen et al., 2015) using LG+F+I+G4 as suggested (BIC criterion) after employing model test implemented in IQ-tree. To milder effects of potential compositional bias in the dataset and long-branch attraction in the 56 universal single copy marker protein trees, the concatenated alignments were re-coded in Dayhoff-4 categories (Schwartz and Dayhoff, 1978; Lartillot et al., 2009; Lartillot et al., 2013) and phylogenetic trees were calculated with PhyloBayesMPI (Lartillot et al., 2013) CAT+GTR in two chains, which both converged with maxdiff=0.09 for the Terrabacteria species tree and maxdiff=0.11 for the detailed species tree. The first 25% of trees in each chain were discarded as burn-in. Phylogenetic tree visualization and annotation was performed with ete3 (Huerta-Cepas et al., 2016).

### Catalytic subunit of NiFe and FeFe hydrogenases and NifH phylogenetic tree

Based on existing annotations target proteins were identified in query proteomes and reference organisms. These proteins were then used to query the NCBI non-redundant (nr) database with diamond blastp (Buchfink et al., 2014). The top 20 hits per query were extracted, merged with queries, de-replicated based on protein accession number and then clustered at 70% and 90% amino acid sequence similarity with cd-hit (Fu et al., 2012). Alignments of cluster representatives were built with MAFFT (v7.294b) (Katoh and Standley, 2013) and maximum-likelihood phylogenetic trees inferred with IQ-tree (multicore v1.5.5) (Nguyen et al., 2015) using the model suggested by the model test feature featured in IQ-tree (based on Bayesian information criterion). Phylogenetic models used were LG+R5 (NifH), LG+R6 (FeFe Hydrogenases) and LG+R8 (NiFe Hydrogenases). To milder effects of long-branch attraction additional phylogenetic trees were constructed with PhyloBayesMPI CAT+GTR (version 1.7) (Lartillot et al., 2013) in two chains, which both converged with maxdiff of 0.18 (NifH tree), 0.16 (FeFeHyd tree), 0.25 (NiFe Hydrogenases). The first approximately 30% of trees were discarded as burn-in. Phylogenetic trees were visualized in ete3 (Huerta-Cepas et al., 2016).

### Catalytic subunit of dimethyl sulfoxide reductase (DMSO) superfamily protein and catalytic subunit I of heme-copper oxygen reductases phylogenetic trees

Each individual protein data set was aligned using Muscle version 3.8.31 (Edgar, 2004a, 2004b) and then manually curated to remove end gaps. Phylogenies were conducted using RAxML-HPC BlackBox (Stamatakis, 2014) as implemented on the CIPRES web server (Miller et al., 2010) under the PROTGAMMA JTT evolutionary model and with the number of bootstraps automatically determined.

### HCO A-family oxygen reductase protein tree

Homologs of CoxA in Marinamargulisbacteria and Saganbacteria genomes were identified from HMM searches. Other homologs were gathered from Shih et al. (2017). Sequences were aligned with MAFFT using the –maxiterate (Katoh et al., 2005). Phylogenetic analysis was performed using RAxML through the CIPRES Science Gateway (Stamatakis, 2014) under the LG model.

### Data availability

New and published genomes included in this study and corresponding gene annotations can be accessed at https://ggkbase.berkeley.edu/Margulis_Sagan_Melaina/organisms (ggKbase is a ‘live’ site, genomes may be updated after publication). Further details are provided in Supplementary Table 1.

## Author contributions

PBMC, IS, FS, BCT, MO and JFB reconstructed and curated the genomes; PBMC conducted the majority of the metabolic analyses, FS, CC, RK, DB, PS, KA and JFB provided input to analyses; FS, CC, PS, and PBMC generated phylogenetic trees; YA, EDB and RS acquired samples; YA, JMS, EDB and RS acquired new sequence information; and EDB, RS and TW contributed single amplified genomes (SAGs). PBMC and JFB wrote the manuscript with input from FS and TW, as well as CC, PS and RK. All authors reviewed the results and approved the manuscript.

## Acknowledgements

Laura Hug and Christopher Brown provided assistance with analyses, Shufei Lei and Kate Lane assisted with bioinformatics and data management, Kelly Wrighton, Kim Handley and Kenneth Hurst Williams provided samples. The research was supported by the Department of Energy (DOE), Office of Science and Office of Biological and Environmental Research (Lawrence Berkeley National Lab; Operated by the University of California, Berkeley). DNA sequencing for the Rifle samples was conducted by the U.S. Department of Energy Joint Genome Institute, a DOE Office of Science User Facility, supported under Contract No. DE-AC02-05CH11231.

Sallie Chisholm, Paul Berube and Steven Biller are acknowledged for their help securing the marine samples. We thank the staff of Bigelow Laboratory Single-Cell Genomics Center for the generation of single-cell data. Marine single amplified genomes were generated and sequenced with the support of NSF grants DEB-1441717 and OCE-1335810, and Simons Foundation grant 510023 (to RS). Teruki Iwatsuki, Kazuki Hayashida, Toshihiro Kato, and Mitsuru Kubota assisted with groundwater sampling at Mizunami Underground Research Laboratory, Japan

Atomic Energy Agency (JAEA). Faecal samples were collected from patients in the clinical PhaseI/II SEASP trial in Bangladesh that was jointly led by Graham George and Ingrid Pickering (University of Saskatchewan), with the assistance of the SEASP team https://clinicaltrials.gov/ct2/show/NCT02377635, and funded by the Canadian Federal Government, through Grand Challenges Canada, Stars in Global Health and by the Global Institute for Water Security.

## Statement of ethics

The faecal samples obtained were part of a clinical PhaseI/II study in rural Bangladesh entitled “Selenium and arsenic pharmacodynamics” (SEASP) run by Graham George (University of Saskatchewan) and funded by the Canadian Federal Government, through a programme entitled Grand Challenges Canada, Stars in Global Health, with additional funds from the Global Institute for Water Security at the University of Saskatchewan. The SEASP trial was approved by the University of Saskatchewan Research Ethics Board (14-284) and the Bangladesh Medical Research Council (940,BMRC/NREC/2010-2013/291). Additional ethics approval was also obtained by UCL (7591/001).

## Competing financial interest

The authors declare no competing financial interests.

## References

Alneberg, J., Bjarnason, B.S., de Bruijn, I., Schirmer, M., Quick, J., Ijaz, U.Z. et al. (2014) Binning metagenomic contigs by coverage and composition. Nat Methods 11: 1144–1146.

Anantharaman, K., Brown, C.T., Hug, L.A., Sharon, I., Castelle, C.J., Probst, A.J. et al. (2016) Thousands of microbial genomes shed light on interconnected biogeochemical processes in an aquifer system. Nat Comm 7: 13219.

Appel, J. (2012) The physiology and functional genomics of cyanobacterial hydrogenases and approaches towards biohydrogen production. In Functional Genomics and Evolution of Photosynthetic Systems. Burnap, R., and Vermaas, W. (eds). Dordrecht: Springer, pp. 357–381.

Baker, B.J., Lazar, C.S., Teske, A.P., and Dick, G.J. (2015) Genomic resolution of linkages in carbon, nitrogen, and sulfur cycling among widespread estuary sediment bacteria. Microbiome 3: 14.

Bankevich, A., Nurk, S., Antipov, D., Gurevich, A.A., Dvorkin, M., Kulikov, A.S. et al. (2012) SPAdes: a new genome assembly algorithm and its applications to single-cell sequencing. J Comput Biol 19: 455–477.

Biegel, E., and Muller, V. (2010) Bacterial Na^+^-translocating ferredoxin:NAD^+^ oxidoreductase. Proc Natl Acad Sci U S A 107: 18138–18142.

Biegel, E., Schmidt, S., and Muller, V. (2009) Genetic, immunological and biochemical evidence for a Rnf complex in the acetogen *Acetobacterium woodii*. Environ Microbiol 11: 1438–1443.

Bothe, H., Schmitz, O., Yates, M.G., and Newton, W.E. (2010) Nitrogen fixation and hydrogen metabolism in cyanobacteria. Microbiol Mol Biol Rev 74: 529–551.

Bowers, R.M., Kyrpides, N.C., Stepanauskas, R., Harmon-Smith, M., Doud, D., Reddy, T.B.K. et al. (2017) Minimum information about a single amplified genome (MISAG) and a metagenome-assembled genome (MIMAG) of bacteria and archaea. Nat Biotechnol 35: 725–731.

Boyd, E.S., Schut, G.J., Adams, M.W., and Peters, J.W. (2014) Hydrogen metabolism and the evolution of biological respiration. Microbe 9: 361–367.

Brown, C.T., Hug, L.A., Thomas, B.C., Sharon, I., Castelle, C.J., Singh, A. et al. (2015) Unusual biology across a group comprising more than 15% of domain Bacteria. Nature 523: 208–211.

Buchfink, B., Xie, C., and Huson, D.H. (2014) Fast and sensitive protein alignment using DIAMOND. Nat Methods 12: 59.

Buckel, W., and Thauer, R.K. (2013) Energy conservation via electron bifurcating ferredoxin reduction and proton/Na^+^ translocating ferredoxin oxidation. Biochim Biophys Acta 1827: 94–113.

Castelle, C.J., Hug, L.A., Wrighton, K.C., Thomas, B.C., Williams, K.H., Wu, D. et al. (2013) Extraordinary phylogenetic diversity and metabolic versatility in aquifer sediment. Nat Commun 4: 2120.

Criscuolo, A., and Gribaldo, S. (2010) BMGE (Block Mapping and Gathering with Entropy): a new software for selection of phylogenetic informative regions from multiple sequence alignments. BMC Evol Biol 10: 210.

Demmer, J.K., Huang, H., Wang, S., Demmer, U., Thauer, R.K., and Ermler, U. (2015) Insights into flavin-based electron bifurcation via the NADH-dependent reduced ferredoxin:NADP oxidoreductase structure. J Biol Chem 290: 21985–21995.

Di Rienzi, S.C., Sharon, I., Wrighton, K.C., Koren, O., Hug, L.A., Thomas, B.C. et al. (2013) The human gut and groundwater harbor non-photosynthetic bacteria belonging to a new candidate phylum sibling to Cyanobacteria. Elife 2: e01102.

Dick, G.J., Andersson, A.F., Baker, B.J., Simmons, S.L., Thomas, B.C., Yelton, A.P., and Banfield, J.F. (2009) Community-wide analysis of microbial genome sequence signatures. Genome biol 10: 1–16.

Edgar, R.C. (2004a) MUSCLE: a multiple sequence alignment method with reduced time and space complexity. BMC Bioinformatics 5: 113.

Edgar, R.C. (2004b) MUSCLE: multiple sequence alignment with high accuracy and high throughput. Nucleic Acids Res 32: 1792–1797.

Eloe-Fadrosh, E.A., Paez-Espino, D., Jarett, J., Dunfield, P.F., Hedlund, B.P., Dekas, A.E. et al. (2016) Global metagenomic survey reveals a new bacterial candidate phylum in geothermal springs. Nat Commun 7: 10476.

Elshahed, M.S., Senko, J.M., Najar, F.Z., Kenton, S.M., Roe, B.A., Dewers, T.A. et al. (2003) Bacterial diversity and sulfur cycling in a mesophilic sulfide-rich spring. Appl Environ Microbiol 69: 5609–5621.

Enright, A.J., Van Dongen, S., and Ouzounis, C.A. (2002) An efficient algorithm for large-scale detection of protein families. Nucleic Acids Res 30: 1575–1584.

Finn, R.D., Clements, J., and Eddy, S.R. (2011) HMMER web server: interactive sequence similarity searching. Nucleic Acids Res 39: W29–37.

Friedrich, T., and Weiss, H. (1997) Modular evolution of the respiratory NADH:ubiquinone oxidoreductase and the origin of its modules. J Theor Biol 187: 529–540.

Friedrich, T., and Scheide, D. (2000) The respiratory complex I of bacteria, archaea and eukarya and its module common with membrane-bound multisubunit hydrogenases. FEBS Lett 479: 1–5.

Fu, L., Niu, B., Zhu, Z., Wu, S., and Li, W. (2012) CD-HIT: accelerated for clustering the next-generation sequencing data. Bioinformatics 28: 3150–3152.

Greening, C., Biswas, A., Carere, C.R., Jackson, C.J., Taylor, M.C., Stott, M.B. et al. (2016) Genomic and metagenomic surveys of hydrogenase distribution indicate H_2_ is a widely utilised energy source for microbial growth and survival. ISME J 10: 761–777.

Hackmann, T.J., and Firkins, J.L. (2015) Electron transport phosphorylation in rumen butyrivibrios: unprecedented ATP yield for glucose fermentation to butyrate. Front Microbiol 6: 622.

Handley, K.M., Wrighton, K.C., Miller, C.S., Wilkins, M.J., Kantor, R.S., Thomas, B.C. et al. (2015) Disturbed subsurface microbial communities follow equivalent trajectories despite different structural starting points. Environ Microbiol 17: 622–636.

Hensel, M., Hinsley, A.P., Nikolaus, T., Sawers, G., and Berks, B.C. (1999) The genetic basis of tetrathionate respiration in *Salmonella typhimurium*. Mol Microbiol 32: 275–287.

Hernsdorf, A.W., Amano, Y., Miyakawa, K., Ise, K., Suzuki, Y., Anantharaman, K. et al. (2017) Potential for microbial H_2_ and metal transformations associated with novel bacteria and archaea in deep terrestrial subsurface sediments. ISME J 11: 1915–1929.

Huerta-Cepas, J., Serra, F., and Bork, P. (2016) ETE 3: Reconstruction, analysis, and visualization of phylogenomic data. Mol Biol Evol 33: 1635–1638.

Hug, L.A., Baker, B.J., Anantharaman, K., Brown, C.T., Probst, A.J., Castelle, C.J. et al. (2016) A new view of the tree of life. Nat Microbiol: 16048.

Hyatt, D., Chen, G.-L., LoCascio, P.F., Land, M.L., Larimer, F.W., and Hauser, L.J. (2010) Prodigal: prokaryotic gene recognition and translation initiation site identification. BMC Bioinformatics 11: 119.

Ino, K., Hernsdorf, A.W., Konno, U., Kouduka, M., Yanagawa, K., Kato, S. et al. (2017) Ecological and genomic profiling of anaerobic methane-oxidizing archaea in a deep granitic environment. ISME J 12: 31.

Jain, C., Rodriguez-R, L.M., Phillippy, A.M., Konstantinidis, K.T., and Aluru, S. (2017) High-throughput ANI analysis of 90K prokaryotic genomes reveals clear species boundaries. bioRxiv: 225342.

Kantor, R.S., Wrighton, K.C., Handley, K.M., Sharon, I., Hug, L.A., Castelle, C.J. et al. (2013) Small genomes and sparse metabolisms of sediment-associated bacteria from four candidate phyla. MBio 4: e00708–00713.

Katoh, K., and Standley, D.M. (2013) MAFFT multiple sequence alignment software version 7: improvements in performance and usability. Mol Biol Evol 30: 772–780.

Katoh, K., Kuma, K., Toh, H., and Miyata, T. (2005) MAFFT version 5: improvement in accuracy of multiple sequence alignment. Nucleic Acids Res 33: 511–518.

Kearse, M., Moir, R., Wilson, A., Stones-Havas, S., Cheung, M., Sturrock, S. et al. (2012) Geneious Basic: an integrated and extendable desktop software platform for the organization and analysis of sequence data. Bioinformatics 28: 1647–1649.

Kim, Y.J., Lee, H.S., Kim, E.S., Bae, S.S., Lim, J.K., Matsumi, R. et al. (2010) Formate-driven growth coupled with H_2_ production. Nature 467: 352–355.

Langmead, B., and Salzberg, S.L. (2012) Fast gapped-read alignment with Bowtie 2. Nat Methods 9: 357–359.

Langmead, B., Trapnell, C., Pop, M., and Salzberg, S.L. (2009) Ultrafast and memory-efficient alignment of short DNA sequences to the human genome. Genome Biol 10.

Lartillot, N., Lepage, T., and Blanquart, S. (2009) PhyloBayes 3: a Bayesian software package for phylogenetic reconstruction and molecular dating. Bioinformatics 25: 2286–2288.

Lartillot, N., Rodrigue, N., Stubbs, D., and Richer, J. (2013) PhyloBayes MPI: phylogenetic reconstruction with infinite mixtures of profiles in a parallel environment. Syst Biol 62: 611–615.

Lea-Smith, D.J., Bombelli, P., Vasudevan, R., and Howe, C.J. (2016) Photosynthetic, respiratory and extracellular electron transport pathways in cyanobacteria. Biochim Biophys Acta 1857: 247–255.

Lolkema, J.S., Chaban, Y., and Boekema, E.J. (2003) Subunit composition, structure, and distribution of bacterial V-type ATPases. J Bioenerg Biomembr 35: 323–335.

Lubner, C.E., Jennings, D.P., Mulder, D.W., Schut, G.J., Zadvornyy, O.A., Hoben, J.P. et al. (2017) Mechanistic insights into energy conservation by flavin-based electron bifurcation. Nat Chem Biol 13: 655–659.

Lücker, S., Wagner, M., Maixner, F., Pelletier, E., Koch, H., Vacherie, B. et al. (2010) A Nitrospira metagenome illuminates the physiology and evolution of globally important nitrite-oxidizing bacteria. Proc Natl Acad Sci U S A 107: 13479–13484.

Luo, G., Ono, S., Beukes, N.J., Wang, D.T., Xie, S., and Summons, R.E. (2016) Rapid oxygenation of Earth’s atmosphere 2.33 billion years ago. Science Advances 2.

Marreiros, B.C., Batista, A.P., Duarte, A.M., and Pereira, M.M. (2013) A missing link between complex I and group 4 membrane-bound [NiFe] hydrogenases. Biochim Biophys Acta 1827: 198–209.

Martinez-Garcia, M., Brazel, D., Poulton, N.J., Swan, B.K., Gomez, M.L., Masland, D. et al. (2012) Unveiling *in situ* interactions between marine protists and bacteria through single cell sequencing. ISME J 6: 703–707.

McDowall, J.S., Murphy, B.J., Haumann, M., Palmer, T., Armstrong, F.A., and Sargent, F. (2014) Bacterial formate hydrogenlyase complex. Proc Natl Acad Sci U S A 111: E3948–3956.

Meuer, J., Kuettner, H.C., Zhang, J.K., Hedderich, R., and Metcalf, W.W. (2002) Genetic analysis of the archaeon *Methanosarcina barkeri Fusaro* reveals a central role for Ech hydrogenase and ferredoxin in methanogenesis and carbon fixation. Proc Natl Acad Sci U S A 99: 5632–5637.

Miller, M.A., Pfeiffer, W., and Schwartz, T. (2010) Creating the CIPRES Science Gateway for inference of large phylogenetic trees. In 2010 Gateway Computing Environments Workshop (GCE), pp. 1–8.

Nguyen, L.-T., Schmidt, H.A., von Haeseler, A., and Minh, B.Q. (2015) IQ-TREE: a fast and effective stochastic algorithm for estimating Maximum-Likelihood phylogenies. Mol Biol Evol 32: 268–274.

Ogata, H., Goto, S., Sato, K., Fujibuchi, W., Bono, H., and Kanehisa, M. (1999) KEGG: kyoto encyclopedia of genes and genomes. Nucleic Acids Res 27: 29–34.

Parks, D.H., Rinke, C., Chuvochina, M., Chaumeil, P.A., Woodcroft, B.J., Evans, P.N. et al. (2017) Recovery of nearly 8,000 metagenome-assembled genomes substantially expands the tree of life. Nat Microbiol 2.

Peng, Y., Leung, H.C.M., Yiu, S.M., and Chin, F.Y.L. (2012) IDBA-UD: a de novo assembler for single-cell and metagenomic sequencing data with highly uneven depth. Bioinformatics 28: 1420–1428.

Peters, J.W., Miller, A.F., Jones, A.K., King, P.W., and Adams, M.W. (2016) Electron bifurcation. Curr Opin Chem Biol 31: 146–152.

Pinske, C., and Sawers, R.G. (2016) Anaerobic formate and hydrogen metabolism. EcoSal Plus 7.

Probst, A.J., Ladd, B., Jarett, J.K., Geller-McGrath, D.E., Sieber, C.M.K., Emerson, J.B. et al. (2018) Differential depth distribution of microbial function and putative symbionts through sediment-hosted aquifers in the deep terrestrial subsurface. Nature Microbiology 3.

Raveh-Sadka, T., Thomas, B.C., Singh, A., Firek, B., Brooks, B., Castelle, C.J. et al. (2015) Gut bacteria are rarely shared by co-hospitalized premature infants, regardless of necrotizing enterocolitis development. Elife 4.

Refojo, P.N., Teixeira, M., and Pereira, M.M. (2012) The Alternative complex III: properties and possible mechanisms for electron transfer and energy conservation. Biochim Biophys Acta 1817: 1852–1859.

Schut, G.J., Bridger, S.L., and Adams, M.W. (2007) Insights into the metabolism of elemental sulfur by the hyperthermophilic archaeon *Pyrococcus furiosus*: characterization of a coenzyme A-dependent NAD(P)H sulfur oxidoreductase. J Bacteriol 189: 4431–4441.

Schwartz, R.M., and Dayhoff, M.O. (1978) Origins of Prokaryotes, Eukaryotes, Mitochondria, and Chloroplasts. Science 199: 395–403.

Sharon, I., Morowitz, M.J., Thomas, B.C., Costello, E.K., Relman, D.A., and Banfield, J.F. (2013) Time series community genomics analysis reveals rapid shifts in bacterial species, strains, and phage during infant gut colonization. Genome Res 23: 111–120.

Shih, P.M., Hemp, J., Ward, L.M., Matzke, N.J., and Fischer, W.W. (2017) Crown group Oxyphotobacteria postdate the rise of oxygen. Geobiology 15: 19–29.

Sieber, C.M.K., Probst, A.J., Sharrar, A., Thomas, B.C., Hess, M., Tringe, S.G., and Banfield, J.F. (2017) Recovery of genomes from metagenomes via a dereplication, aggregation, and scoring strategy. bioRxiv: 107789.

Soo, R.M., Hemp, J., Parks, D.H., Fischer, W.W., and Hugenholtz, P. (2017) On the origins of oxygenic photosynthesis and aerobic respiration in Cyanobacteria. Science 355: 1436–1440.

Soo, R.M., Skennerton, C.T., Sekiguchi, Y., Imelfort, M., Paech, S.J., Dennis, P.G. et al. (2014) An expanded genomic representation of the phylum Cyanobacteria. Genome Biol Evol 6: 1031–1045.

Stamatakis, A. (2014) RAxML version 8: a tool for phylogenetic analysis and post-analysis of large phylogenies. Bioinformatics 30: 1312–1313.

Stepanauskas, R., Fergusson, E.A., Brown, J., Poulton, N.J., Tupper, B., Labonte, J.M. et al. (2017) Improved genome recovery and integrated cell-size analyses of individual uncultured microbial cells and viral particles. Nat Commun 8: 84.

Suzek, B.E., Huang, H., McGarvey, P., Mazumder, R., and Wu, C.H. (2007) UniRef: comprehensive and non-redundant UniProt reference clusters. Bioinformatics 23: 1282–1288.

Tatusov, R.L., Galperin, M.Y., Natale, D.A., and Koonin, E.V. (2000) The COG database: a tool for genome-scale analysis of protein functions and evolution. Nucleic Acids Res 28: 33–36.

Ultsch, A., and Mörchen, F. (2005) ESOM-Maps: tools for clustering, visualization, and classification with Emergent SOM: Univ.

Vermaas, W.F.J. (2001) Photosynthesis and respiration in Cyanobacteria. In eLS: John Wiley & Sons, Ltd.

Vignais, P.M., and Billoud, B. (2007) Occurrence, classification, and biological function of hydrogenases: an overview. Chem Rev 107: 4206–4272.

Welte, C., Kratzer, C., and Deppenmeier, U. (2010) Involvement of Ech hydrogenase in energy conservation of *Methanosarcina mazei*. FEBS J 277: 3396–3403.

Woyke, T., Xie, G., Copeland, A., González, J.M., Han, C., Kiss, H. et al. (2009) Assembling the marine metagenome, one cell at a time. PLoS ONE 4: e5299.

Wrighton, K.C., Thomas, B.C., Sharon, I., Miller, C.S., Castelle, C.J., VerBerkmoes, N.C. et al. (2012) Fermentation, hydrogen, and sulfur metabolism in multiple uncultivated bacterial phyla. Science 337: 1661–1665.

Wu, Y.-W., Simmons, B.A., and Singer, S.W. (2016) MaxBin 2.0: an automated binning algorithm to recover genomes from multiple metagenomic datasets. Bioinformatics 32: 605–607.

Yu, F.B., Blainey, P.C., Schulz, F., Woyke, T., Horowitz, M.A., and Quake, S.R. (2017) Microfluidic-based mini-metagenomics enables discovery of novel microbial lineages from complex environmental samples. Elife 6.

Zerbino, D.R., and Birney, E. (2008) Velvet: algorithms for de novo short read assembly using de Bruijn graphs. Genome Res 18: 821–829.

